# Comparison of Genomic Selection Models for Exploring Predictive Ability of Complex Traits in Breeding Programs

**DOI:** 10.1101/2021.04.15.440015

**Authors:** Lance F. Merrick, Arron H. Carter

**Author notes:** **Correspondence:** Dr. Arron H. Carter.

## Abstract

Traits with a complex unknown genetic architecture are common in breeding programs. However, they pose a challenge for selection due to a combination of complex environmental and pleiotropic effects that impede the ability to create mapping populations to characterize the trait’s genetic basis. One such trait, seedling emergence of wheat (*Triticum aestivum* L.) from deep planting, presents a unique opportunity to explore the best method to use and implement GS models to predict a complex trait. 17 GS models were compared using two training populations, consisting of 473 genotypes from a diverse association mapping panel (DP) phenotyped from 2015-2019 and the other training population consisting of 643 breeding lines phenotyped in 2015 and 2020 in Lind, WA with 40,368 markers. There were only a few significant differences between GS models, with support vector machines reaching the highest accuracy of 0.56 in a single breeding line trial using cross-validations. However, the consistent moderate accuracy of cBLUP and other parametric models indicates no need to implement computationally demanding non-parametric models for complex traits. There was an increase in accuracy using cross-validations from 0.40 to 0.41 and independent validations from 0.10 to 0.17 using diversity panels lines to breeding lines. The environmental effects of complex traits can be overcome by combining years of the same populations. Overall, our study showed that breeders can accurately predict and implement GS for a complex trait by using parametric models within their own breeding programs with increased accuracy as they combine training populations over the years.

## Introduction

Plant breeding programs select and improve a wide variety of traits, each with its own challenges and genetic control, that a breeder must overcome to increase genetic gains. In addition to environmental effects from abiotic and biotic factors, agronomic traits interact with each other causing pleiotropic effects resulting in an unknown complex genetic architecture controlled by many quantitative trait loci (QTL; Luo et al. 2017). The identification of QTLs either through linkage mapping or genome-wide association studies (GWAS) allows marker-assisted selection (MAS) and introgression to efficiently select and deploy specific markers in a population (Lande and Thompson 1990). MAS is effective for large-effect alleles with useful markers but has proven to be challenging to use with complex quantitative traits controlled by multiple, minor effect alleles (Bernardo 2008).

These complex traits are common in breeding programs but pose a challenge for selection and significant genetic gain. Challenges due to complexity, combined with the difficulty in phenotyping, can impede the ability to create useful mapping populations for linkage mapping or GWAS to characterize the genetic basis of traits using QTLs. Complex traits controlled by many minor QTLs can result in inconsistent estimated QTL effects (Bernardo 2008). Inconsistent estimations create a need for large mapping populations in each breeding program and environment. Genome-wide association mapping is a newer alternative to biparental linkage mapping but is also limited by complex traits and relies on identifying only significant loci that are common in the germplasm (Bernardo 2014). Even with newer models that can dissect complex traits, GWAS studies rely on statistical thresholds to determine the significance of a marker and identify QTLs.

Previous methods to deal with the limitations of identifying markers for quantitative traits were developed, such as a multi-marker MAS system, but it was difficult to identify and account for all the effects (Bernardo 2008, 2010). Therefore, Meuwissen et al. (2001) discussed the idea of simultaneously estimating all markers regardless of “significance” and thereby capturing all effects. Meuwissen coined this method “Genomic Selection” (GS) (Meuwissen et al. 2001; Lorenz et al. 2011). GS increases genetic gains and reduces cycle time for complex agronomic traits difficult to phenotype and analyze. With the advent of high-throughput genotyping, it is now feasible to develop and implement GS models for complex traits that breeders deal with on a routine basis, such as traits with unknown genetic architecture. The accuracy of GS is affected by an array of factors, such as the prediction model used.

Different GS models can be used to account for genetic effects and genetic architecture. GS models are created using parametric, non-parametric, and semi-parametric approaches. The most common methods are parametric models that include ridge regression best linear unbiased prediction (rrBLUP) and Bayesian approaches. Parametric or linear models assume a linear relationship between the phenotype and genotype, which accounts for only additive effects (Pérez-Rodríguez et al. 2012). Nonlinear models (semi-parametric and non-parametric) such as support vector machines (SVM), can account for both additive and non-additive genetic effects (e.i. dominance and epistasis) and can improve accuracy for traits with complex genetic basis (Desta and Ortiz 2014; Varona et al. 2018).

The accuracy of the different models varies with the genetic architecture of a trait due to their assumptions and treatment of marker effects (Larkin et al. 2019). Genomic best linear unbiased predictions (gBLUP) have been independent of the number of QTLs controlling the trait whereas Bayesian models show increased accuracy with fewer QTLs controlling the trait (Daetwyler et al. 2010; Wang et al. 2018). In smaller training sets, rrBLUP outperformed Bayesian models, presumably due to less allelic variation needed to predict small effect QTL (Heffner et al. 2011a). Reproducing kernel Hilbert spaces (RKHS) and random forest (RF) models have improved prediction accuracies in maize for complex traits with epistasis over Bayesian and best linear unbiased predictor (BLUP) based models in wheat (*Triticum aestivum* L.), maize (*Zea mays* L.), and barley (*Hordeum vulgare* L.) (Dudley and Johnson 2009; Pérez-Rodríguez et al. 2012; Heslot et al. 2013; Jia 2017; Shikha et al. 2017).

Therefore, the best model differs between the trait and crop of interest, making plant breeders test multiple models to find the model with the best fit and highest accuracy (Larkin et al. 2019). By comparing models for the highest accuracy, a breeder can determine and exploit the genetic architecture for increased genetic gains. Seedling emergence in deep-sown winter wheat exhibits an unknown complex genetic architecture with a significant environmental effect and provides an opportunity for researchers to employ GS methods to select for complex traits.

Seedling emergence is dependent on deep sowing in Mediterranean climates such as the Pacific Northwest to reach stored moisture when precipitation is below 150 to 300 mm annually (Schillinger et al. 1998; Mohan et al. 2013). In low-precipitation dryland areas, fast-emerging cultivars are the most desirable because rain events before emergence create soil crusting and decreased seedling emergence. In deep planting cropping systems, stand establishment and emergence are vital factors affecting grain yield and can reduce grain yields by 35 to 40% (Schillinger et al. 1998). Decreased emergence is caused by the wheat seedling’s inability to penetrate the soil surface (Schillinger et al. 1998) or by the inability of the seed to germinate under dry soil conditions. Seedling emergence in deep-sown winter wheat depends on the first true leaf’s ability to protroude through the coleoptile and reach the soil surface before a crusting event (Schillinger 2011). Therefore, emergence is affected by the speed of emergence, force exerted, and lifting capacity of the first leaf (Arndt 1965; Schillinger et al. 2017; Lutcher et al. 2019). These studies showed that the genetic basis of seedling emergence is a complex trait affected by many factors and is dependent on the environment to display variation (Schillinger et al. 2017; Lutcher et al. 2019).

The inability for mapping studies, the complex nature, and environmental dependency create a need for plant breeders and geneticists to investigate GS to select and improve complex traits (Bernardo 2008). This study presents empirical research to assess the predictive ability and genetic architecture of a complex trait using GS without the need for mapping studies. This study’s objective was to compare GS models for seedling emergence using different types of training populations within and across years to account for genetic relatedness and environmental variation. The results of this study helped determine the best method for implementing GS for a complex trait such as seedling emergence in deep-sown winter wheat.

## Material and Methods

### Phenotypic Data

The Washington State University winter wheat breeding program takes emergence notes every fall to select for strong emerging lines in low rainfall (∼150 mm annual precipitation), deep furrow trial locations. Two training populations were used to compare models. The first training population consists of a diverse association mapping panel (DP) that was evaluated in 2015, 2017, 2018, and 2019 in Lind, WA (Table 1). The second training population consists of unreplicated trials of F_3:5_ and doubled-haploid (DH) breeding lines (BL) that were evaluated in 2015 and 2020 (Table 1). In 2016, no data was collected for either population due to significant soil crusting that delayed seedling emergence of all lines. There was not enough seedling emergence variation for adequate phenotyping in the BL trials from 2017-2019. The two training populations comprise two scenarios in a breeding program. The DP represents a diverse panel of lines evaluated each year. Whereas, the BL population represents closely related lines from a single breeding program comprised of different types of trials that are combined in a way to imitate real-life scenarios in a breeding program. The BL trials are combined based on availability from year to year.

**Table 1.**
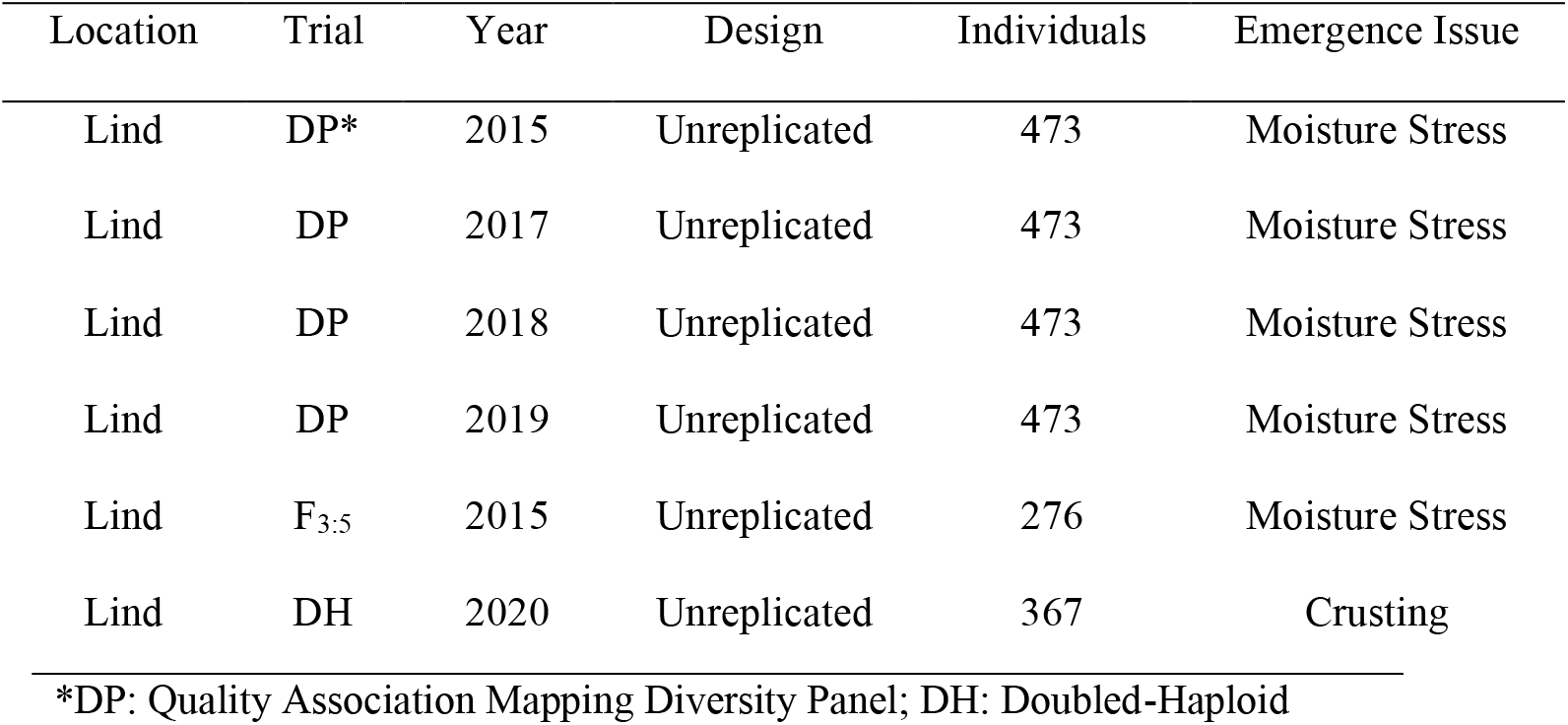
Study Populations for Seedling Emergence.

| Location | Trial | Year | Design | Individuals | Emergence Issue |
| --- | --- | --- | --- | --- | --- |
| Lind | DP* | 2015 | Unreplicated | 473 | Moisture Stress |
| Lind | DP | 2017 | Unreplicated | 473 | Moisture Stress |
| Lind | DP | 2018 | Unreplicated | 473 | Moisture Stress |
| Lind | DP | 2019 | Unreplicated | 473 | Moisture Stress |
| Lind | F<sub>3:5</sub> | 2015 | Unreplicated | 276 | Moisture Stress |
| Lind | DH | 2020 | Unreplicated | 367 | Crusting |

\*DP: Quality Association Mapping Diversity Panel; DH: Doubled-Haploid

For each trial, seedling emergence was measured. Seedling emergence was visually assessed and recorded as a percentage of the total plot emerged six weeks after planting. Table 1 summarizes location, year, field design, number of genotyped individuals, and emergence issue for each trial where seedling emergence was recorded.

A two-step adjusted means method was used in which a linear model was used to adjust means within and across environments. A mixed linear model was then used to calculate genomic estimated breeding values (GEBVs; Ward et al. 2019). Adjusted means from the emergence data collected in the unreplicated trials were adjusted using residuals calculated for the unreplicated genotypes in individual environments and across environments using the modified augmented complete block design model (ACBD; Federer 1956; Goldman 2019). The modified model excludes entry effect to prevent running out of degrees of freedom for best linear unbiased estimates (BLUEs) in a fixed effect model or from overfitting BLUPs in a random effect model in our cross-validation scheme. The model in a single environment is as follows:

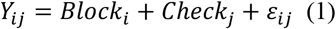

Where *Y*_*ij*_ is the trait of interest; *Block*_*i*_ is the fixed effect of the ith block; *Check*_*j*_ is the fixed effect of the jth replicated check cultivar; and *ε*_*ij*_ are the residual errors. For adjusted means across environments, the model is as follows:

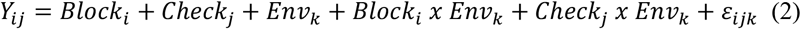

Where *Y*_*ij*_ is the trait of interest; *Block*_*i*_ is the fixed effect of the ith block; *Check*_*j*_ is the fixed effect of the jth replicated check cultivar; *Env*_*k*_ is the fixed effect of the kth environment; and *ε*_*ijk*_ are the residual errors.

BLUPs for the GWAS models and heritability were calculated for each trial and across trials using a mixed linear model for the full augmented randomized complete block design in a single environment and is as follows:

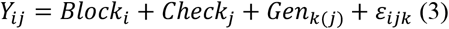

Where *Y*_*ij*_ is the trait of interest; *Block*_*i*_ is the random effect of the ith block; *Check*_*j*_ is the fixed effect of the jth replicated check cultivar; *Gen*_*k*(*j*)_ is the genotype k in the jth Check; and *ε*_*ij*_ are the residual errors. For the full model (3) across environments is as follows:

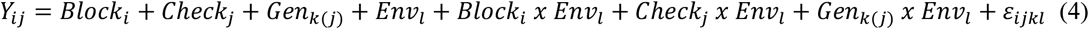

Where *Y*_*ij*_ is the trait of interest; *Block*_*i*_ is the random effect of the ith block; *Check*_*j*_ is the fixed effect of the jth replicated check cultivar; *Gen*_*k*(*j*)_ is the genotype k in the jth Check; *Env*_*l*_ is the random effect of the lth environment; and *ε*_*ijkl*_ are the residual errors. Heritability on a genotype-difference basis for broad-sense heritability is then calculated from the variance components using model (3) for both individual environments and across environments using the formula (Cullis et al. 2006):

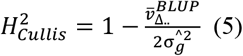

where 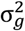, and 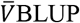 is the genotype variance and mean-variance of the BLUPs (Cullis et al. 2006). Trial evaluation and significant differences were evaluated using the coefficient of variation, and analysis of variance (ANOVA) in individual and across trials.

### Genotypic Data

Lines were genotyped using genotyped-by-sequencing (GBS; Elshire et al. 2011) through the North Carolina State Genomics Sciences laboratory in Raleigh, NC, using the restriction enzymes *Msp*I and *Pst*I (Poland et al. 2012). Sequences were aligned to the Chinese Spring International Wheat Genome Sequencing Consortium (IWGSC) RefSeq v1.0 (Appels et al. 2018), using the Burrows-Wheeler Aligner (BWA) 0.7.17 (Li and Durbin 2009). Genetic markers with more than 20% missing data, minor allele frequency of less than 5%, and those that were monomorphic were removed. Markers were then imputed using Beagle version 5.0 (Browning et al. 2018). A total of 40,368 single-nucleotide polymorphism (SNP) markers remained. The 473 DP lines were phenotyped for seedling emergence over four years (2015, 2017, 2018, and 2019). 643 BL lines were phenotyped for seedling emergence over two years (2015 and 2020). Principal component (PC) analysis biplots were developed using the markers combined over both the DP and BL using the function “prcomp” in R (R Core Team 2018).

### Genomic Selection Models

#### Parametric Models

Genomic selection models have to incorporate both genotypic and phenotypic data that need to estimate a large number of marker effects compared to phenotypic data. This creates a dilemma with the small p, large n problem which results in a lack of degrees of freedom and low statistical power. In order for parametric models to overcome this, popular genomic selection models use various methods such as shrinkage procedures. The parametric models used include rrBLUP, gBLUP, compressed BLUP (cBLUP), and super BLUP (sBLUP), referred to together as BLUP models. The Bayesian prediction models used were the BayesA, BayesB, BayesCπ, Bayesian LASSO (BayesL), and Bayesian Ridge Regression (BayesRR). rrBLUP and gBLUP were implemented using the package ‘rrBLUP’, with gBLUP using a marker-based relationship matrix (Endelman 2011), whereas the Bayesian models were implemented using the package ‘BGLR’ (Pérez and de los Campos 2014). All models implemented in BGLR used a burn-in rate of 2,000 and 5,000 iterations. Both cBLUP and sBLUP were implemented using GAPIT with three principal components as fixed effects (Tang et al. 2016; Wang et al. 2018).

The parametric models integrated three principal components to account for population structure and follow the basic mixed linear model that treat the effects of markers as random effects and principal components as fixed effects:

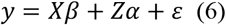

Where y is a vector of phenotypes; *β* is a vector of non-genetic fixed effects; X is an incidence matrix for the fixed effects *β*; *α* is a vector of random regression coefficients of all the marker effects; Z is an n x k genotypic matrix for markers; and *ε* is a vector of residuals. Principal components were calculated using ‘prcomp’ in R (R Core Team 2018).

#### Semi-Parametric Models

The semi-parametric models compared are the RKHS “single-kernel” model and the “Multikernel” (kernel averaging; MK) methods described in Perez and de los Campos (2017). RKHS models are an alternative to parametric models for capturing complex interactions that account for both additive and epistatic effects. The Bayesian RKHS regression model described in Perez and de los Campos (2017) is:

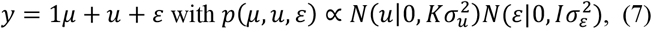

Where K={*K*(*x*_*j*_, *x*_*i*_)} is an (n x n)-matrix with observations are evaluations of the RK at pairs of points. In the “single-kernel” or “average” RKHS, a gaussian kernel derived from the squared-Euclidean distance between genotypes is:

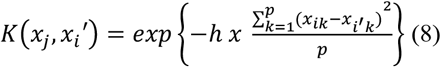

In “Multikernel” RKHS, kernel averaging consists of defining a sequence of kernels and fitting a multikernel model with random effects as kernels, which follow the model as described in Perez and de los Campos (2017):

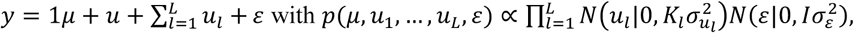

Where *K*_*l*_ is the RK evaluated at the lth value of the parameter in the sequence {*h*_1_, …, *h*_*L*_}. The regression function follows the model:

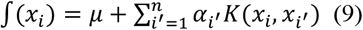

Where *x*_*i*_ = (*x*_*i*1_, …, *x*_*ip*_) ′ and *x*_*i*_ = (*x*_*i*′1_, …, *x*_*i*′*p*_) ′ are input vectors of marker genotypes in individuals i and *i*′; *α*_*i*′_ are regression coefficients; and K(*x*_*i*_, *x*_*i*′_) = exp (−*h*│|*x*_*i*_ − *x*_*i*′_|│^2^) is the reproducing kernel with a gaussian radial basis function (RBF), where h is a bandwidth parameter and │|*x*_*i*_ − *x*_*i*′_ |│ is the Euclidean norm between each pair of input vectors. The kernel averaging for selecting optimal values of h within a set of candidate values is implemented using a Bayesian approach (Pérez-Rodríguez et al. 2012). Both models were implemented in “BGLR” and fitted with fixed effects composed of principal components, and a burn-in rate of 2,000 and 5,000 iterations (Pérez and de los Campos 2014).

#### Non-Parametric Models

The non-parametric models compared were RF, and SVM with an RBF. SVMs and RF are supervised machine-learning method for both classification and regression. They have been applied to overcome the large p, small n mentioned previously. SVM regression maps samples from a predictor space to a high-dimensional feature space using a non-linear kernel function and then completes linear regression in the feature space (Jannink et al. 2010). RF predictors consist of a collection of tree-regression predictors, where each tree in the group is developed using bootstrapped samples of the training set. Each tree predicts the response, and the group of tree regressions predicts the response as an average of the individual tree predictions (Jannink et al. 2010).

RF and SVM prediction models were executed using the package ‘caret’ (Kuhn 2008). All packages were implemented in R (R Core Team 2018). Model tuning was completed using ten replications of ten-fold CV. Three different forms of feature selection were used to implement the non-parametric models. The first type of feature selection used was the sampling of 3,000 random markers with the RF and SVM indicated with RF_M and SVM_M. The 3,000 markers were used to reduce the number of predictor variables while still maintaining enough markers to capture the majority of linkage disequilibrium. In a previous study, it has been shown that over 1000 SNPs have been required to capture the majority of variation (Sandhu et al. 2020). The second type of feature selection was completed using principal components to reduce the dimensionality of the genotypic data, and the number of principal components for each training population was equivalent to the number of individuals phenotyped for each cross-validation training fold (Song et al. 2010). RF and SVM are indicated by RF_PC and SVM_PV, respectively. The last feature selection was implemented using a reduced genotype matrix of 26,253 SNPs that was composed of markers that were binned together based on a threshold LD value of 0.80 (Ward et al. 2019). The reduced genotype matrix was computed using JMP genomics version 9 (SAS Institute, Inc 2011). After the genotype matrix was reduced, a random sample of 3,000 markers was used, and the RF and SVM models are indicated by RF_LD and SVM_LD, respectively. In all, there were six models used for non-parametric model comparisons.

### Prediction Accuracy and Cross-Validation Scheme

Prediction accuracy for the GS was reported using Pearson correlation coefficients between GEBVs and their respective adjusted means using the function “cor” in R (R Core Team 2018). GS for emergence was assessed using cross-validations and independent validations using the DP and BL trials as the two training populations. The different populations were used to compare the DP to the BL for prediction purposes and compare whether it would be beneficial to grow independent populations outside of the breeding program for GS purposes. GS models were conducted with five-fold cross-validation by including 80% of the samples in the training population and predicting the GEBVs of the remaining 20% (Lozada and Carter 2019). One replicate consists of the five model iterations, where the population is split into five different groups. This was completed 50 times. Independent validations were then performed using the combined years training population to predict individual years in the other training population. The genomic selection scheme is shown in figure 1.

**Figure 1.**
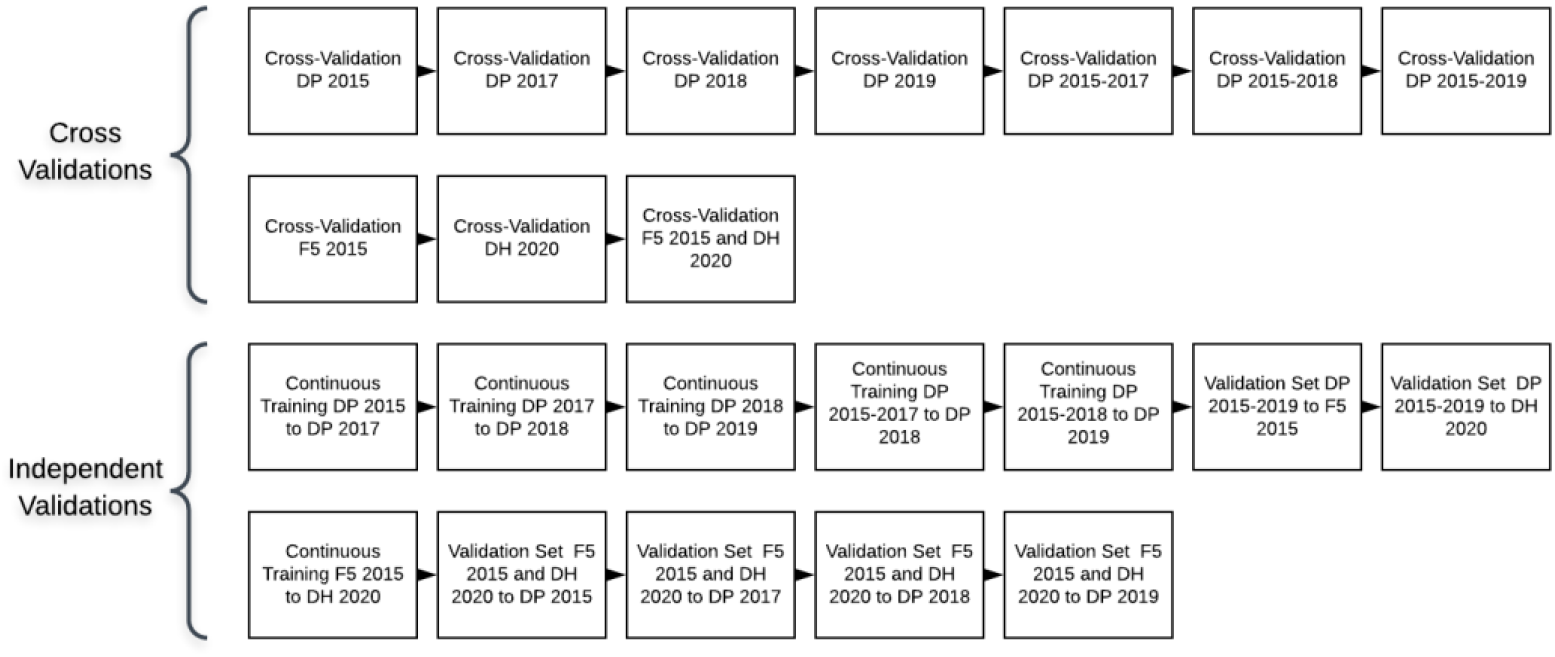
Genomic selection scheme for the cross-validation and independent validations using the diversity panel (DP) and breeding line (BL) training populations consisting of the F_3:5_ and doubled-haploid (DH) trials from 2015 to 2020.

The training populations were evaluated for cross-validations on a yearly/trial basis and over combined years/trials. We assessed each trial independently using cross-validations. We then created prediction models starting with the earliest trial and then a new model with the addition of each subsequent trial to evaluate genotype-by-environment interaction and continuous training of a prediction model. The independent validations were first conducted using continuous training. For example, the first trial, such as DP 2015, is used to predict the subsequent trial, DP 2017. Then we used the previous years combined to predict the next year such as combine DP 2015-2017 and predict DP 2018. The validation sets were then completed using the combined trials in one training population to predict individual trials in the other training population, such as DP 2015-2019 to predict the F_3:5_ breeding lines trial. All GS models and scenarios were analyzed using Washington State University’s Kamiak high performance computing cluster (Kamiak 2021).

### Model Comparison

Model, scenario, and training populations comparisons were evaluated by using a Tukey’s honestly significant difference (HSD) test implemented in the “agricolae” package in R (R Core Team 2018; de Mendiburu and de Mendiburu 2019). The comparison of models was then plotted for visual comparison using “ggplot2” in R (Wickham 2011; R Core Team 2018).

## Results

### Phenotypic Data

Seedling emergence varies widely depending on the year due to the need for specific environmental conditions to promote variation between genotypes. The heritability in individual trials was higher than in combined trials in both the DP and the BL. The highest heritability in a single trial was 0.88 in the DP in 2018, and the highest heritability in combined trials was 0.66 in the BL for the combination of BL trials (Table 2). The trials with minimum values of zero have a larger standard deviation, which indicates a wider range of seedling emergence and environmental pressure and phenotypic variation for selection purposes.

**Table 2.**
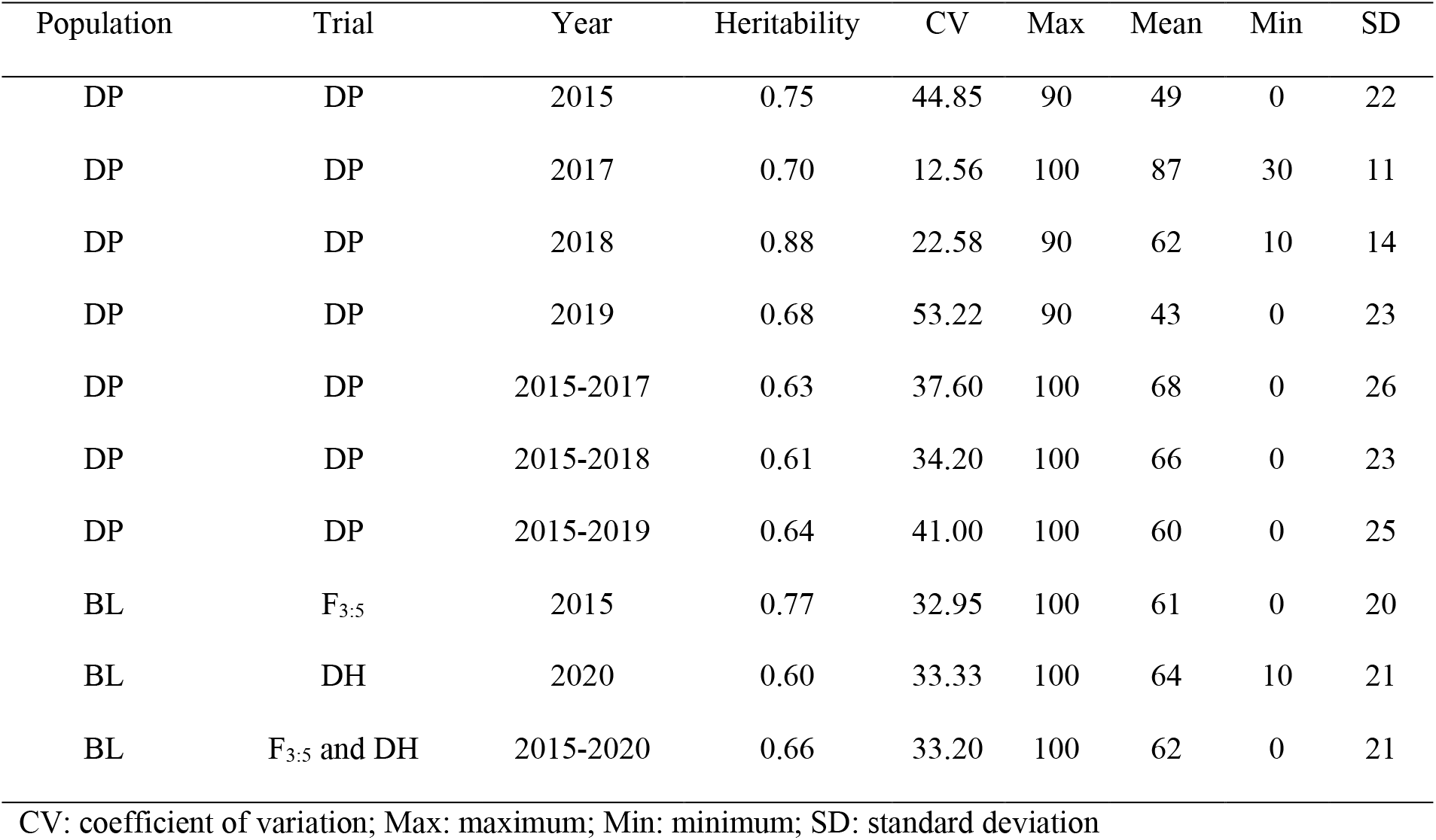
Deep-sowing seedling emergence heritability and trial statistics for the diversity panel (DP) training population and breeding line (BL) training population phenotyped from 2015 to 2019 and 2015 to 2020, respectively.

| Population | Trial | Year | Heritability | CV | Max | Mean | Min | SD |
| --- | --- | --- | --- | --- | --- | --- | --- | --- |
| DP | DP | 2015 | 0.75 | 44.85 | 90 | 49 | 0 | 22 |
| DP | DP | 2017 | 0.70 | 12.56 | 100 | 87 | 30 | 11 |
| DP | DP | 2018 | 0.88 | 22.58 | 90 | 62 | 10 | 14 |
| DP | DP | 2019 | 0.68 | 53.22 | 90 | 43 | 0 | 23 |
| DP | DP | 2015-2017 | 0.63 | 37.60 | 100 | 68 | 0 | 26 |
| DP | DP | 2015-2018 | 0.61 | 34.20 | 100 | 66 | 0 | 23 |
| DP | DP | 2015-2019 | 0.64 | 41.00 | 100 | 60 | 0 | 25 |
| BL | F<sub>3:5</sub> | 2015 | 0.77 | 32.95 | 100 | 61 | 0 | 20 |
| BL | DH | 2020 | 0.60 | 33.33 | 100 | 64 | 10 | 21 |
| BL | F<sub>3:5</sub> and DH | 2015-2020 | 0.66 | 33.20 | 100 | 62 | 0 | 21 |

CV: coefficient of variation; Max: maximum; Min: minimum; SD: standard deviation

Since the accuracy of GS and the variation for seedling emergence is dependent on the environmental effect, it is important to examine the variance components for the trials. Genotypic variances were significant for the DP in 2015, 2017, and for all of the combined trials (Table S1). The environmental effect was not significant in any of the combined trials, with the genotype-by-environment interaction (GEI) effect only significant in the DP over the combined trials of 2015-2018 and 2015-2019 (Table S1).

The majority of GS applications are in single environments due to the lack of predictive power to make selections across multiple environments. Therefore, it is important to understand the correlations between training populations and environments for predictive purposes. The DP trials are significantly positively correlated to each other with the exception of three scenarios: DP 2015 to DP 2018; DP 2017 to DP 2018; and DP 2017 to DP 2019 (Table S2). The BL trials, F_3:5_ 2015 and DH 2020, were negatively correlated to each other. The F_3:5_ 2015 trial was significantly correlated to DP 2017, and DH 2020 was significantly correlated to DP 2018. The combined trials were significantly correlated with the single trials that are included in them.

### Genotypic Data

The principal component biplot using the SNP markers for the DP and BL training population accounted for 15% of the total genetic variation (Figure 2). PC1 explained 8% of the variation, and PC2 explained 6% of the variation. The biplot revealed two evident subpopulations, with the majority of BL lines clustering with some of the DP and the other cluster consisting mainly of DP lines. Both the F_3:5_ and DH breeding line trials are grouped together.

**Figure 2.**
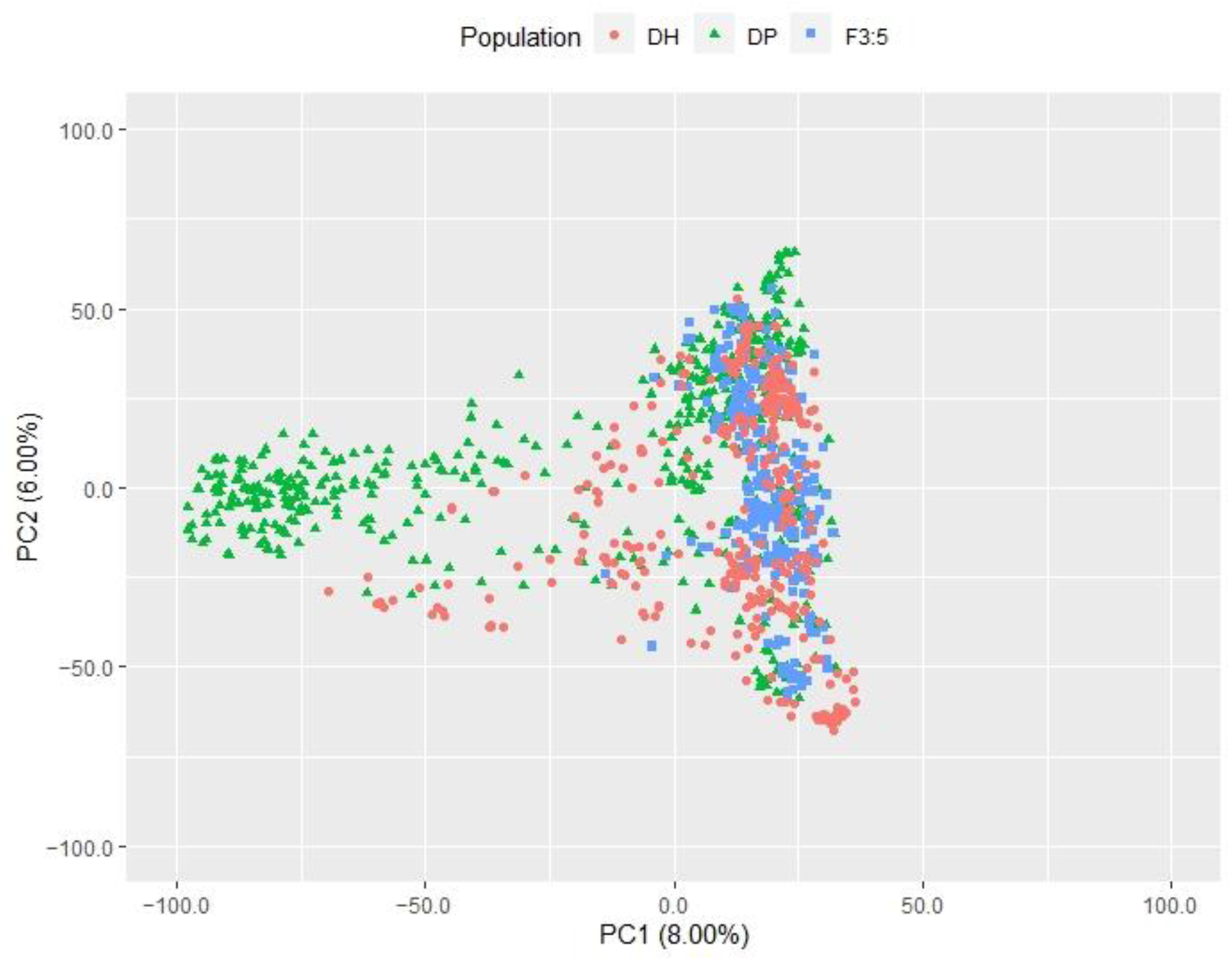
Principal Component (PC) biplot of genetic markers from the diversity panel (DP) and breeding line (BL) training populations consisting of the F_3:5_ and doubled-haploid (DH) trials.

### Genomic Selection Model Comparison

Multiple comparisons were completed for the GS models for each training population in the DP and BL. Overall, there were only a few statistical differences between GS models in most training populations. Depending on the DP training population, the average accuracy across individual and combined training populations ranged from 0.31 to 0.42 for all models (Table S3). The models that performed consistently worse than the rest of the models were sBLUP, RF_PC, and SVM_PC. Parametric models consistently had the highest mean accuracies. The Bayesian models had very similar accuracy within each training population, and there were only a few significant differences between the Bayesian models and the BLUP models. The semi-parametric models, RKHS and MK, had statistically similar accuracy to the parametric models. In a year with more variation between significant differences such as 2019, the parametric model cBLUP had a statistically higher mean than the other parametric models (Figure 3). The RF models using markers (RF_M and RF_LD) and SVM_LD also had statistically similar accuracy to cBLUP. By combining the trials, the accuracy of the models increased. The combined trials had the highest mean accuracy for most of the models, with the combined trials of DP 2015-2018 having the highest mean accuracy for every model (Figure 4). RF_M reached the highest accuracy of 0.52.

**Figure 3.**
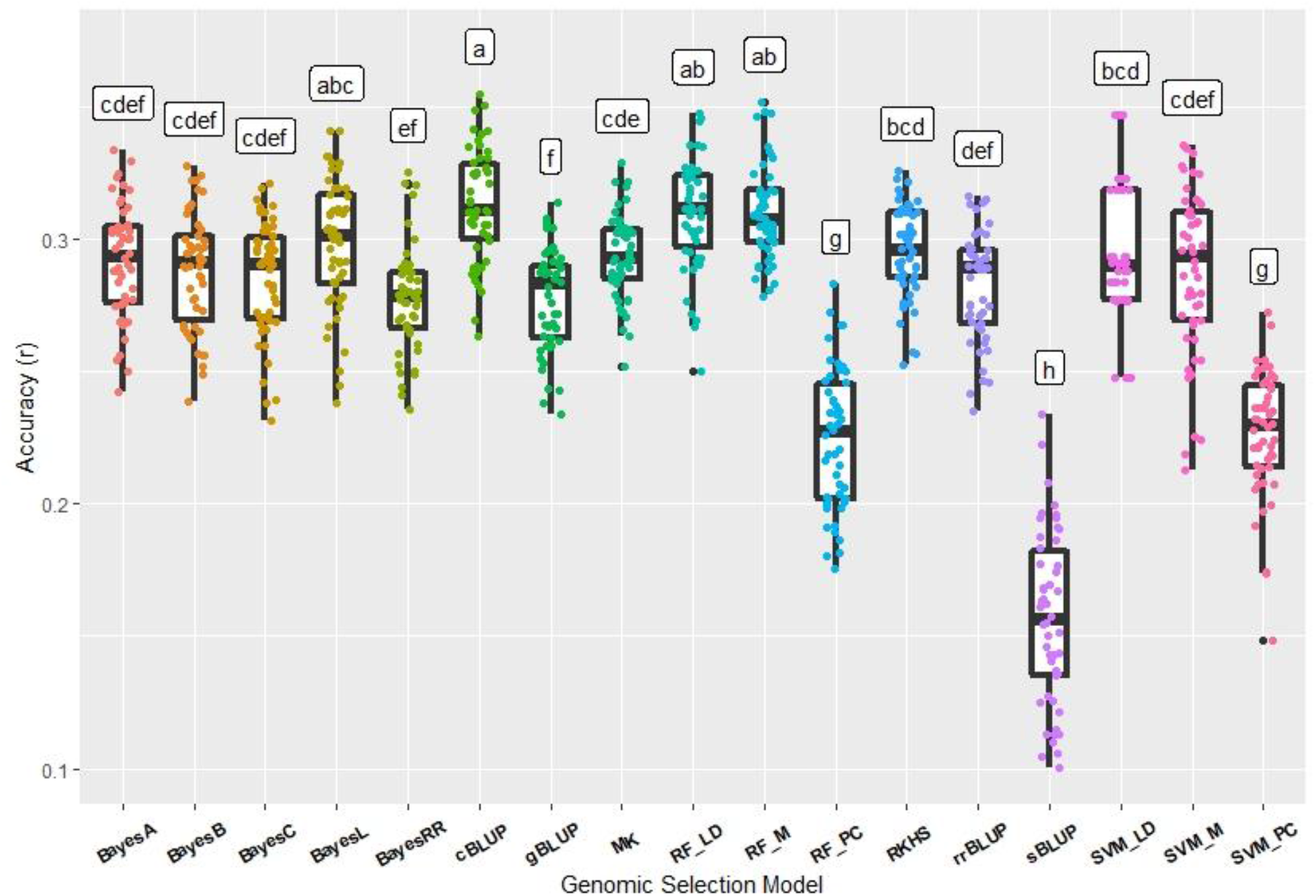
Comparison of genomic selection model accuracy and pairwise comparisons using cross-validation for deep-sowing seedling emergence for Pacific Northwest winter wheat diversity panel (DP) lines phenotyped in 2019 in Lind, WA. Models labeled with the same letter are not significantly different (P-value =0.05).

**Figure 4.**
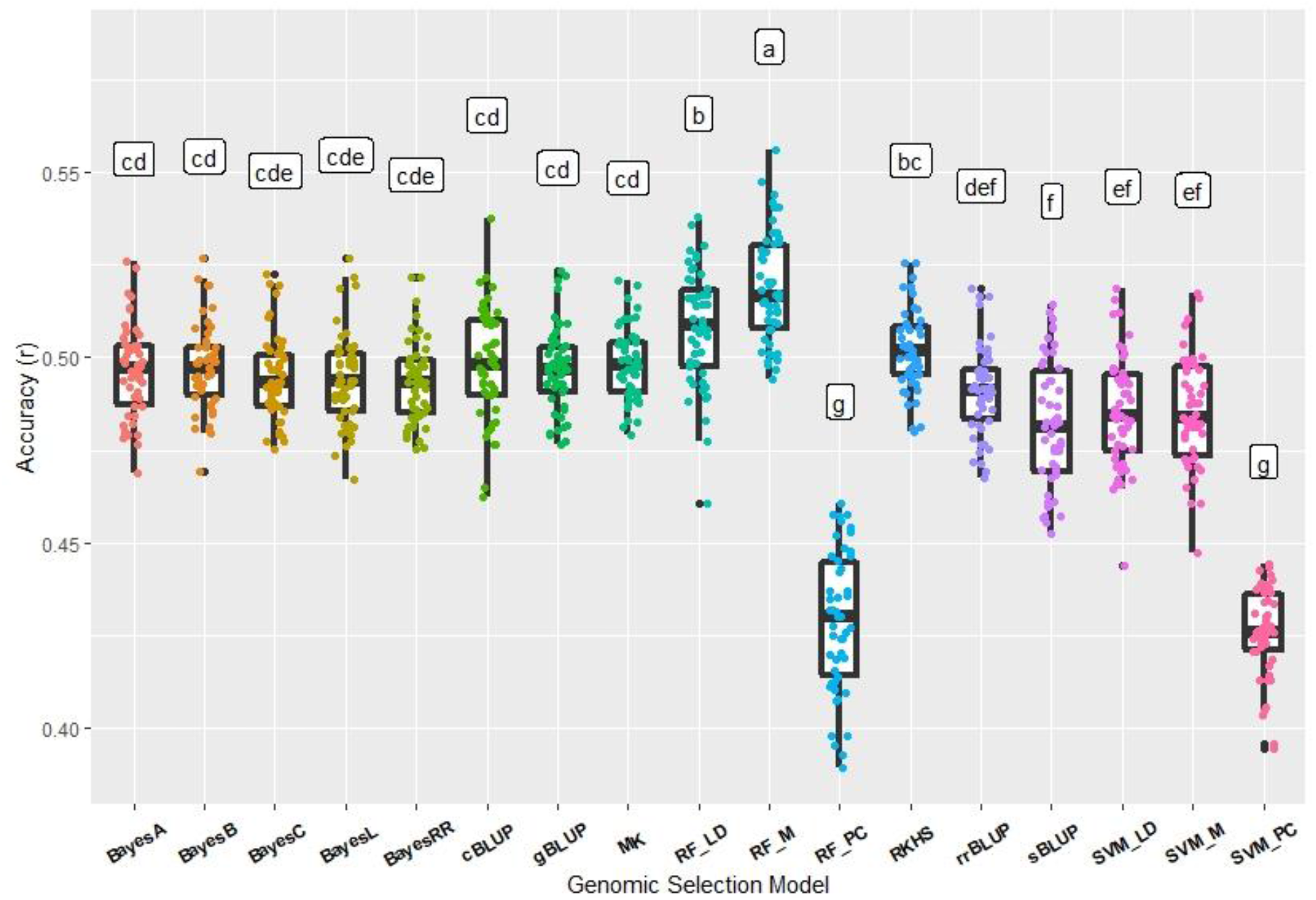
Comparison of genomic selection model accuracy and pairwise comparisons using cross-validation for deep-sowing seedling emergence for Pacific Northwest winter wheat diversity panel (DP) lines phenotyped from 2015-2018 in Lind, WA. Models labeled with the same letter are not significantly different (P-value =0.05).

The model accuracy between the DP and BL training populations were comparable. There were only a few statistical differences between GS models using the BL training populations. The highest accuracies over all training populations reached 0.56 for SVM_M and 0.55 for cBLUP, MK, RKHS, RF_LD, RF_M, and SVM_LD (Table S4). Similar to the DP training populations, the lowest accuracy models were sBLUP, RF_PC, and SVM_PC. In the combined trial, F_3:5_ and DH, the semi-parametric models, RKHS and MK, had the highest mean accuracies of 0.44 (Figure 5). The parametric BLUP models rrBLUP, gBLUP, and cBLUP had statistically similar accuracy in the combined trial. In the BL training populations, the non-parametric models had the highest accuracy in a single trial as seen in the DP training populations, but did not have the highest accuracy in the combined trials.

**Figure 5.**
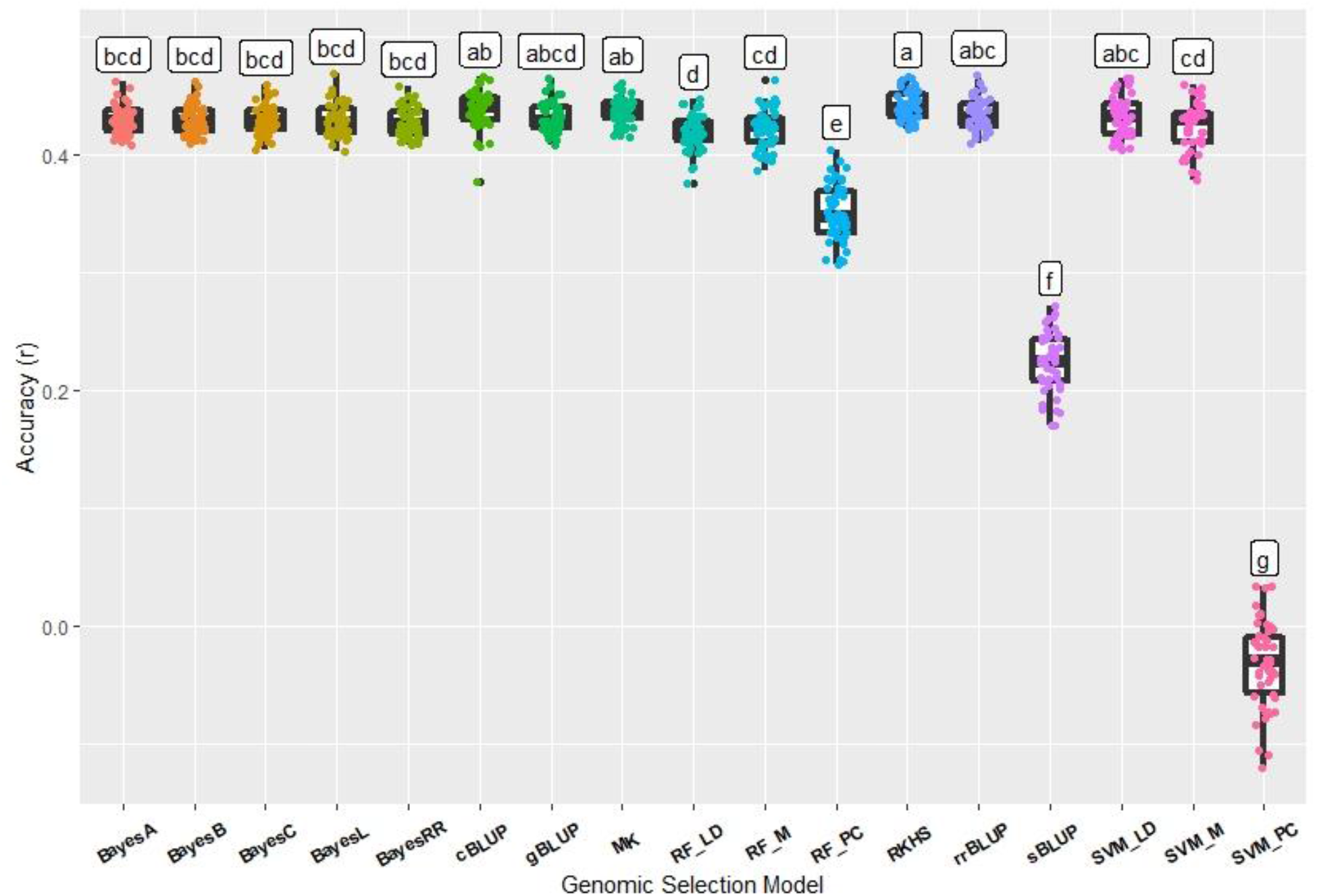
Comparison of genomic selection model accuracy and pairwise comparisons using cross-validation for deep-sowing seedling emergence for Pacific Northwest winter wheat breeding lines phenotyped across 2015 F_3:5_ and 2020 DH in Lind, WA. Models labeled with the same letter are not significantly different (P-value =0.05).

### Continuous Training and Validation Sets

Independent validation sets displayed more statistical differences between models. Continuous training schemes included predicting a trial or year using the previous year simulating scenarios in breeding programs. The accuracy was lower than the cross-validation accuracy and neared zero for a few scenarios (Table S5). Using the DP 2015 to predict the DP 2017 resulted in the highest mean accuracies in the DP continuous training scenarios. The RF_PC model resulted in the highest mean accuracy of 0.22 (Figure 6). In the DP 2015 to DP 2017 continuous training scenario, the parametric and non-parametric models had statistically higher mean accuracies than the SVM and other RF models. But, there were only a few statistical differences between the models within each model type. There were very few significant differences in statistical accuracy between any of the models in the single BL continuous training scenario (Table S6). All of the models except for sBLUP had negative accuracies for the F_3:5_ to DH continuous training independent validation. cBLUP reached similar accuracies to the other independent validations but was negative (−0.28).

**Figure 6.**
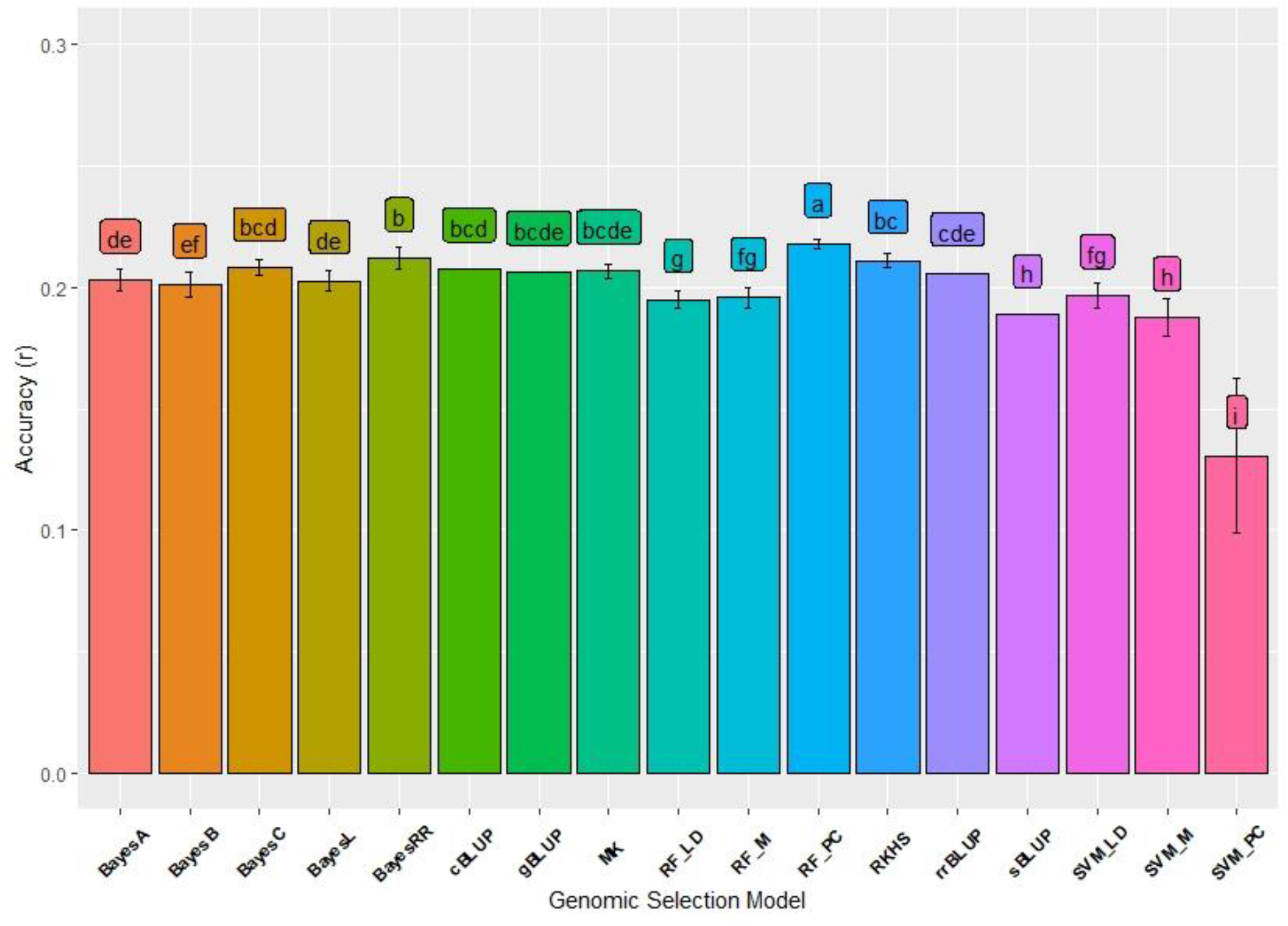
Comparison of genomic selection model accuracy and pairwise comparisons using continuous training for deep-sowing seedling emergence for Pacific Northwest winter wheat diversity panel (DP) lines phenotyped in 2015 in Lind, WA to predict the DP lines phenotyped in 2017 in Lind, WA. Models labeled with the same letter are not significantly different (P-value =0.05)

Validation sets were completed using the DP to predict the BL and vice versa.; the two validation sets scenarios for the DP in which the combined trials of DP 2015-2019 predicting the BL trials resulted in low accuracy with both positive and negative accuracy depending on the model (Table S5). Overall, the parametric and semi-parametric models had the highest mean accuracies in the DP validation sets. The BL validation set scenarios in which the combined BL trials predicted individual DP trials resulted in higher accuracy than the DP validation sets (Table S6). The prediction models had the same accuracy regardless of the DP trial, with cBLUP and RF_M having the highest mean accuracy of 0.32 (Figure 7). The RF_LD was the only model with any difference in accuracy between the BL validation sets.

**Figure 7.**
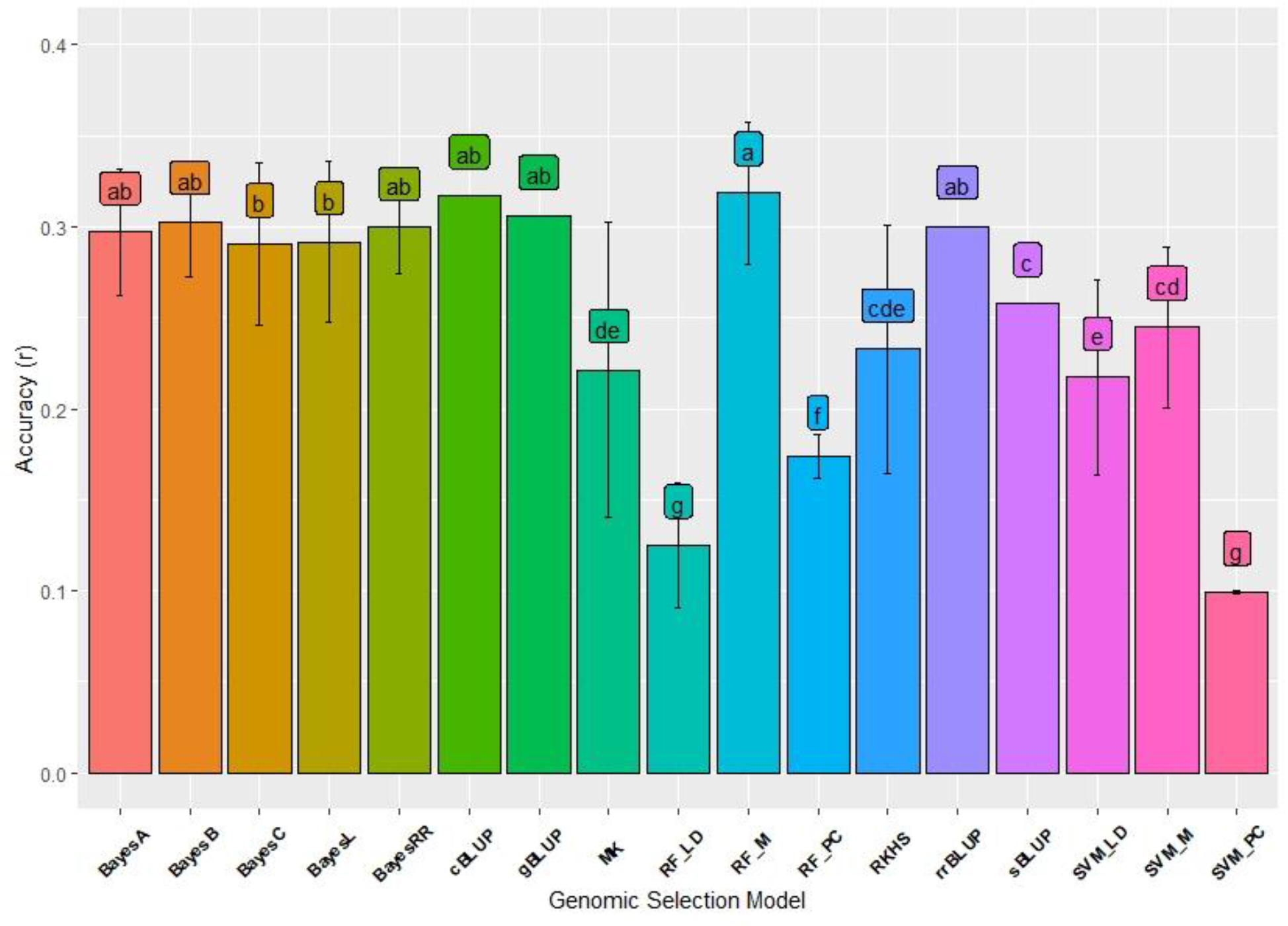
Comparison of genomic selection model accuracy and pairwise comparisons using independent validation for deep-sowing seedling emergence for Pacific Northwest winter wheat breeding lines phenotyped in 2015 and 2020 in Lind, WA to predict the DP lines phenotyped in 2017 in Lind, WA. Models labeled with the same letter are not significantly different (P-value =0.05).

### Training Population Comparison

Using pairwise comparisons, we compared the training populations and types of prediction scenarios to better understand the predictive ability within and across trials. As seen in the comparison of GS models, there is a statistical increase in prediction accuracy over combined trials in the DP training populations; however, this is not seen in the BL training populations (Table S7; Figure 8). The cross-validations of DP 2015-2018 and the 2015 F3:5 trials reached the highest mean accuracy of 0.49. The BL cross-validations had the widest range of accuracies with the SVM models reaching low to negative accuracies.

**Figure 8.**
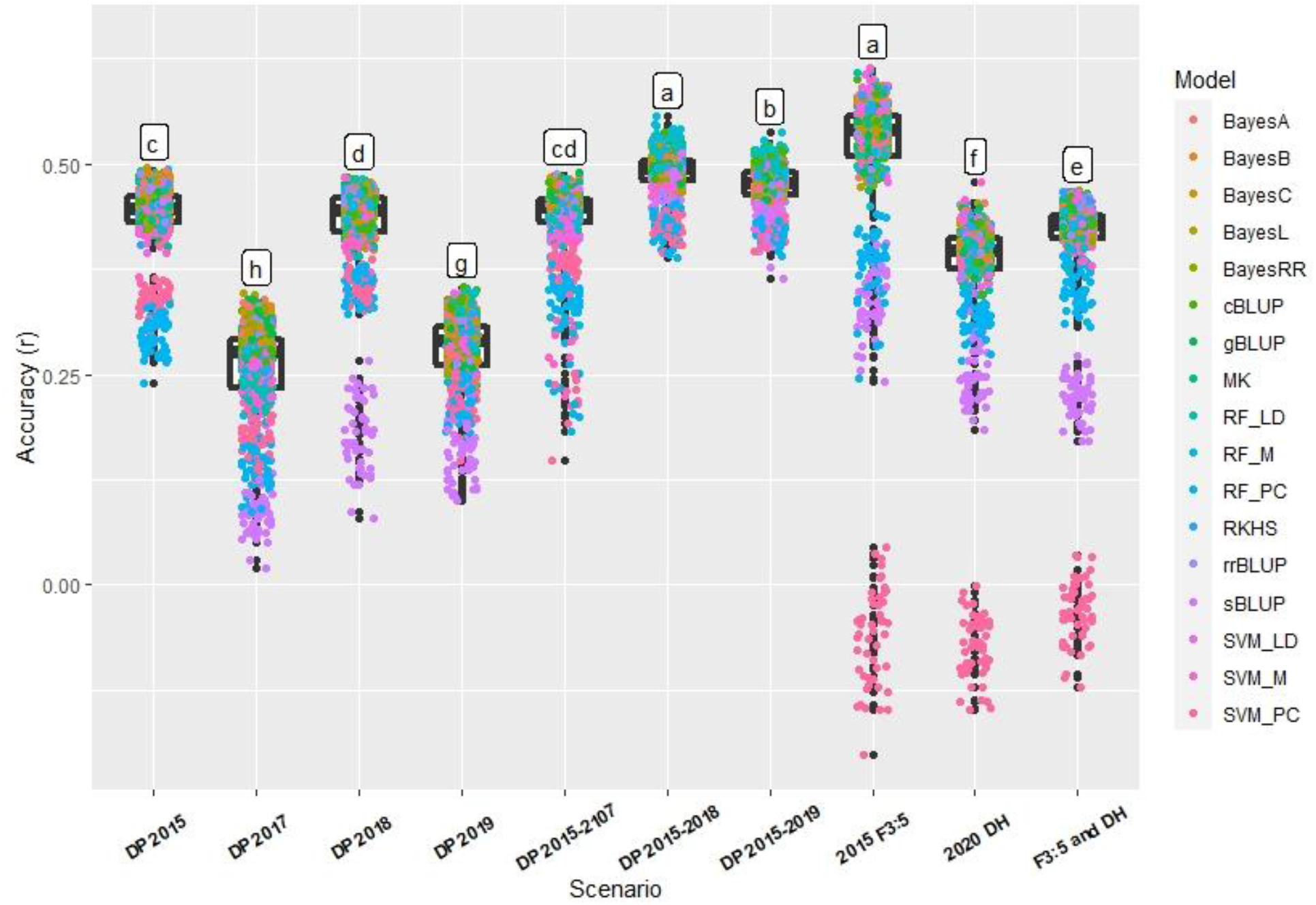
Comparison of accuracy and pairwise comparisons between cross-validation scenarios across all genomic selection models for deep-sowing seedling emergence for Pacific Northwest winter wheat diversity panel (DP) lines and breeding lines phenotyped across 2015 to 2020 in Lind, WA. Models labeled with the same letter are not significantly different (P-value =0.05).

The continuous training scenarios vary in mean accuracy, with DP 2015 to DP 2017 having the highest mean accuracy within all DP independent validations (Table S7; Figure 9). The continuous training scenario for the BL training population had the lowest mean accuracy with −0.14. The validation sets for the BL to DP had the highest mean accuracy with very similar results for all four DP trials. The two validation sets for the DP training population had very low mean accuracy than the breeding line training population. All of the validations set in the BL training populations had higher accuracy than all of the DP independent validations and the BL continuous training scenario.

**Figure 9.**
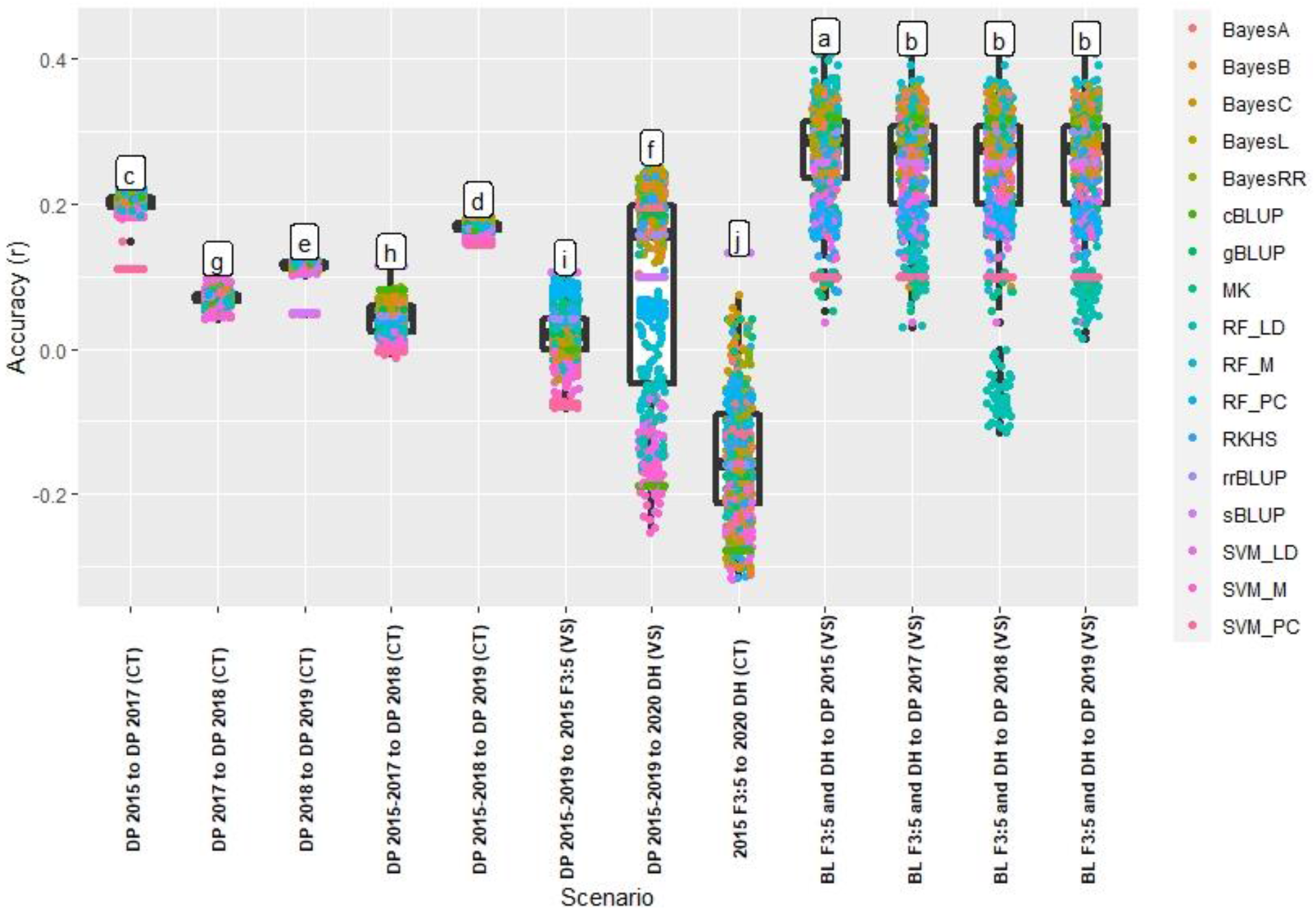
Comparison of accuracy and pairwise comparisons between the two independent validation scenarios, continuous training (CT) and validation sets (VS). The accuracy was compared across all genomic selection models for deep-sowing seedling emergence for Pacific Northwest winter wheat diversity panel (DP) lines and breeding lines phenotyped across 2015 to 2020 in Lind, WA. Models labeled with the same letter are not significantly different (P-value =0.05).

When we compared each scheme over all models for each training population, the cross-validations had statistically higher accuracy for both the DP and BL (Table 5). As shown previously, the DP had a higher statistical accuracy for continuous training than the BL, but the BL had a higher statistical accuracy than the DP in validation sets. Over both training populations, the validation sets had a higher statistical accuracy than the continuous training scenarios. When combining the continuous training and validation sets, the BL training populations had statistically higher accuracy in cross-validations, independent validations, and overall.

**Table 5.** Comparison of genomic selection schemes and populations for deep-sowing seedling emergence for cross-validations and both types of independent validations, continuous training and validation sets, in a Pacific Northwest winter wheat diversity panel and breeding lines phenotyped in 2015-2020 in Lind, WA.

| Scheme | Diversity Panel | Breeding Lines | Overall |
| --- | --- | --- | --- |
| Cross Validations | 0.4a | 0.41a | 0.4a |
| Continuous Training | 0.12b | -0.14c | 0.08c |
| Validation Sets | 0.05c | 0.25b | 0.19b |
| Population | Cross Validations | Independent Validations | Overall |
| Diversity Panel | 0.4b | 0.1b | 0.25b |
| Breeding Lines | 0.41a | 0.17a | 0.26a |

Models labeled with the same letter are not significantly different (P-value =0.05)

## Discussion

### Complex Traits and Seedling Emergence

Complex traits are quantitative in nature and are affected by many small-effect QTLs, the environment, and the interaction between the QTLs and the environment (Holland 2007). Complex traits are common in breeding programs but pose challenges to characterize the genetic architecture. The challenges that impede the understanding of complex traits are the inability to statistically detect and map minor effect QTLs, accurately understand GEI, and account for pleiotropic effects (Luo et al. 2017). However, just because a trait is complex or has an unknown genetic architecture does not mean that breeding programs cannot select or improve these traits. However, this gain can be slow and tedious such as the less than 1% annual increase of yield (Lopes et al. 2012).

Seedling emergence was previously shown to be largely dependent on coleoptile length (Allan et al. 1962; Sunderman 1964; Chastain et al. 1995; Schillinger et al. 1998; Botwright et al. 2001; Schillinger 2011). With the recent study conducted by Mohan et al. (2013), coleoptile length was shown to only account for 28% of the variation of seedling emergence in modern genotypes. The need to understand the genetic architecture of seedling emergence has increased but has been impeded by the multitude of factors that effect emergence. It is not feasible to create linkage mapping for the first leaf force, lifting capacity, speed of emergence, and coleoptile length and diameter especially since emergence is dependent on both moisture and the lack of crusting events to promote adequate phenotypic variation. Therefore, GS will be key for future selection and improvement of these types of complex, pleiotropic traits. In this study, we demonstrated the ability to predict complex traits while deducing the most accurate model, population, and implementation strategy.

### Genomic Selection Models

Genome-wide association mapping is a newer alternative to biparental analyses, but is also limited by complex traits and relies on identifying only significant loci that are common in the germplasm (Bernardo 2014). The genetic architecture of complex traits and the resources needed for genetic mapping and mapping populations make GS an ideal tool for plant breeders (Larkin et al. 2019). GS presents a unique opportunity to account and select for seedling emergence based on genome-wide markers without the need to identify significant markers or fully characterize the genetic architecture.

The difference in GS model accuracy has been shown to be due to genetic architecture (Daetwyler et al. 2010). Genetic architecture is dependent on the number of QTLs and the heritability of the trait (Wang et al. 2018). The accuracy of GS models varies with the genetic architecture of a trait due to their assumptions and treatment of marker effects. In our study there was a lack of significant differences between the common parametric Bayesian models, rrBLUP and gBLUP which support results presented previously in predicting complex traits (Daetwyler et al. 2010; Heffner et al. 2011b; Heslot et al. 2012; Lorenz et al. 2012; Rolling et al. 2020). In early simulation studies for GS, BayesB had higher accuracy than gBLUP, but in more empirical studies the results are very similar (Meuwissen et al. 2001; Habier et al. 2007; Daetwyler et al. 2010). gBLUP had a constant accuracy dependent on the number of individuals in a training population and heritability and assumes a polygenic architecture. Whereas, BayesB was also dependent on the number of loci controlling the trait, with a decreasing accuracy as the number of loci increased (Daetwyler et al. 2010). The two types of models were similar with a large number of controlling loci with gBLUP occasionally showing a higher accuracy (Daetwyler et al. 2010). gBLUP and rrBLUP have been shown to be independent of the number of QTL associated with the phenotypic traits. Both rrBLUP and gBLUP treat markers homogeneously with a common variance and has the ability to accurately predict complex quantitative traits with many minor effect QTLs (Endelman 2011). Whereas BayesB and other Bayesian models (BayesA, BayesC, and BayesL) are sensitive to the number of QTL and are more effective for predicting qualitative traits (Meuwissen et al. 2001; Habier et al. 2007; Daetwyler et al. 2010; Wang et al. 2015).

As previously mentioned, in addition to the effects of genetic architecture, heritability can affect prediction accuracy (Desta and Ortiz 2014). Highly heritable traits are positively correlated with higher prediction accuracy specifically, but complex traits such as yield have been shown to have lower heritability (Lozada et al. 2020). We obtained moderate heritabilities for seedling emergence with higher heritabilities than yield (Lozada et al. 2019, 2020; Lozada and Carter 2020). This may be due to the Cullis heritability model accurately calculating heritability, but also displays why the GS models obtained higher prediction accuracies than grain yield for GS models on these same training populations used in prior GS studies (Lozada et al. 2019, 2020).

Two other genomic selection models, cBLUP and sBLUP, were developed for specific genetic architecture and heritability (Wang et al. 2018). Our results show that sBLUP consistently had a lower mean accuracy compared to most models, and cBLUP was one of the top-performing GS models. Even though there were very few significant differences between the bayesian and BLUP models, cBLUP consistently had higher accuracy in accurately modeling the complex nature of seedling emergence. cBLUP is more accurate for low heritable complex traits, and sBLUP is more accurate for high heritable qualitative traits (Wang et al. 2018). sBLUP has been found superior to most prediction models for a trait controlled by a few QTLs with high heritability similar to Bayesian models and concurs why it consistently had poor predictive ability. cBLUP has been shown to be superior to most models except when the trait is controlled by few QTLs, and is similar or superior to gBLUP, which can also be seen in our study (Wang et al. 2018).

The GS models we have discussed already have all been parametric models. Parametric models only model additive effects, but complex traits have been shown to be affected by non-additive effects (Balestre et al. 2010; Wellmann and Bennewitz 2012; Denis and Bouvet 2013). Therefore, semi-parametric models and non-parametric models have been used to model both additive and non-additive effects of complex traits. Common semi-parametric models include kernel regression, RKHS. RKHS has been used for various regression and classification problems and introduced for genomic selection with Gianola et al. (2006). Both RKHS models, MK and RKHS, showed similar accuracy to both the parametric and non-parametric models in our study. RKHS models have been shown to have similar to higher accuracy for complex traits than parametric models, which have also been shown in our results (Pérez-Rodríguez et al. 2012; Juliana et al. 2017). In our study, the non-parametric models, SVM and RF, proved to have the highest mean accuracy in the DP population and in a single trial in the BL (0.56). The RF and SVM models did display higher accuracy than the parametric models in some of the scenarios in our study, which supports results seen in previous studies (González-Camacho et al. 2018; Azodi et al. 2019; Đorđević et al. 2019).

In our study, we also compared different methods for implementing the non-parametric models. There was no significant difference in the majority of trials between the regular genotype matrix sample (RF_M and SVM_M) and the LD reduced genotype matrix (RF_LD and SVM_LD). This demonstrates that our original genotype matrix may have had redundant markers that were in LD with each other. For both the regular genotype matrix-based and LD reduced genotype matrix-based models we used random sampling to reduce computing time. Random sampling of markers has been noted to be a poor feature selection and not recommended (Pérez-Enciso and Zingaretti 2019). However, the accuracy of the models using random sampling was not statistically different from the parametric and non-parametric models and even had higher accuracy in some scenarios. We also implemented PCs to reduce the dimensionality and improve computing time for our non-parametric models. The PC reduced models (RF_PC and SVM_PC) proved to have the lowest mean accuracy over the majority of populations and scenarios. Using PCs to reduce the dimensionality of our predictor variables reduced the accuracy in the majority of scenarios, indicating either overfitting of the model or a loss of variation due to reducing dimensionality.

The lack of statistical difference between the models in the majority of scenarios indicates little to no advantage in the model by accounting for non-additive effects (semi-parametric and non-parametric) compared to accounting for only additive effects (parametric). The similarity between the parametric models and semi and non-parametric models indicates that either the non-additive effects are not present or small enough that they are negligible (Riedelsheimer et al. 2012). Either way, the Bayesian models, semi-parametric, and non-parametric models are more computationally demanding than the BLUP models with no significant advantage in accuracy. Therefore, rrBLUP or cBLUP is recommended for implementing GS for complex traits with unknown genetic architecture.

### Effect of Prediction scenario and the Environment

It is important to understand and compare prediction scenarios due to their effect on accuracy and the potential implementation of GS into breeding programs. Cross-validations displayed significantly higher mean accuracy over both models and training populations than both types of independent validations (continuous training and validation sets). The increase in accuracy using cross-validation results from the same environment and population and has been shown in previous studies (Lozada and Carter 2019; Đorđević et al. 2019; Haile et al. 2020). We even see an increase in prediction accuracy in cross-validations using multiple trials and environments when we use the same population. The statistically higher accuracy using cross-validations over the independent validations shows that cross-validations may be a more reasonable approach in integrating GS for complex traits.

However, the importance of testing and using independent validations cannot be overstated as it displays more realistic expectations and practical uses of genomic selection in breeding programs (Lozada and Carter 2019). The accuracy of independent validations being lower than cross-validations can be contributed to by the larger differences between populations as compared to within populations along with GEI effect between trials (Michel et al. 2016; Huang et al. 2018; Lozada and Carter 2019, 2020; Haile et al. 2020). Independent validations using continuous training or validation sets of GS models is important for GS in plant breeding programs (Lozada and Carter 2019). As new generations are introduced into the breeding program, genetic differences grow with the introduction of new parents into the crossing blocks. With the retraining of the models comes the combination of multiple environments and GEI. For a trait, such as seedling emergence, the environment plays an important role in creating conducive conditions to promote genotypic variation adequate for phenotypic selection. The environmental requirement of seedling emergence screening is why GS can play a helpful role in selection. Therefore, understanding the environmental effects on GS accuracy is crucial for adequately implementing GS for complex traits such as deep-sown seedling emergence in winter wheat.

In our results, we used a two-step adjusted means model to account for GEI for GS purposes outlined in Ward et al. (2019), because accounting for GEI on complex traits is important. Following the two-step adjusted means model, we implemented a linear model within and across environments, then used a second mixed linear model to calculate GEBVs. The two-step adjusted means model was shown to perform similarly to more complicated GEI incorporated models (Ward et al. 2019). We see an increase in GS accuracy as we combine trials in the cross-validations, presumably due to accounting for the GEI. This increase in accuracy with the inclusion of GEI has been shown in previous studies (Jarquín et al. 2014; Crossa et al. 2014; Haile et al. 2020). There was an increase in accuracy in the cross-validations for the DP. There was also an increase in the BL combined trial over the F_3:5_ trial but not over the DH trial. This discrepancy was presumably due to the large difference between cross-validation accuracy within each trial and the negative accuracy of predicting the DH 2020 trial from the F_3:5_ 2015 trial. We see the DP increase in accuracy because it consisted of the same genotypes from year to year, and by combining the trials, we were able to replicate genotypes and partition the GEI from the genotype effect and accurately predict the genotype effect.

There are generally two environmental scenarios that cause issues with deep-sowing seedling emergence and allows for field screening. The first is when there is not enough moisture in the soil to allow the first true leaf to reach the surface. The second is when rainfall occurs after planting but before emergence, causing soil crusting in which the first true leaf cannot penetrate (Schillinger 2011). For example, in 2015 through 2019, the soil was very dry, so the variation in emergence was due to the ability of the genotype to overcome the first scenario and emerge in the face of moisture stress, but in 2020 the variation in emergence was due to crusting and the ability of either fast emerging genotypes or the ability to penetrate the crust.

The environmental dependence creates two problems to account for and overcome to implement GS. First, when there is not enough environmental effect to display genetic variation, the GS models fail to model the differences due to genotypes adequately. The lack of adequate screening conditions is similar to phenotypic selection problems and is easy to overcome by not including field trials such as the BL trials from 2016 to 2019 in our training populations. Secondly, since the cause of poor seedling emergence can be attributed to both genotypic and different environmental causes, GS scenarios using individual trials and independent validations may not be able to account for the cause of poor emergence from one year to the next. However, by combining our trials, we are able to increase our prediction accuracy in both our cross-validation and independent validation scenarios. The BL training populations are the only trials that combine both causes of seedling emergence for training our GS models. This may explain why we see moderate to similar accuracy in our validation set comparable to our cross-validations when we predict the DP using the combined BL. This difference between causes of seedling emergence is also one reason we see reduced and negative accuracy when we predict one BL trial using the other BL trial. Another example of this effect is why continuous training scenarios result in higher accuracy for the DP but not BL because the DP predicts a single cause of poor seedling emergence. Therefore, it is important to create training populations that include both causes of poor seedling emergence and try to account for as many of the factors affecting seedling emergence as possible.

### Training Populations

The composition of the training populations is a major factor in the prediction accuracy of GS models (Lorenz et al. 2011; Desta and Ortiz 2014; Larkin et al. 2019). It is important to understand and optimize the training population for the GS models and their implementation into a breeding program. We used both a DP and BL training population to determine whether a more diverse panel of genotypes compared to breeding lines from the same breeding program would be more beneficial in using GS for a complex trait. Training population composition is important due to the effect of population structure on GS model accuracy (Asoro et al. 2011). We evaluated the population composition of our training populations by using PCs. The PCs displayed an overlap from the DP and BL training populations. However, the majority of the DP clustered away from the BL, indicating a difference in genetic relatedness, and two distinct subpopulations. We used PCs as fixed effects in our parametric and semi-parametric GS models to account for and remove bias due to the presence of underlying population structure in our training populations.

For complex traits, GS can exploit the genetic relatedness of the training populations for predicting genotypes (Habier et al. 2007; Mirdita et al. 2015). The genetic similarity between training populations and the validation sets can significantly affect prediction accuracy (Habier et al. 2007; Bernardo 2020). The closer related the genotypes are, the fewer recombination events, thus preserving marker linkage disequilibrium, and less genetic variation needed to be accounted for by the models. The difference between the DP and BL training populations is that the Washington State University breeding lines have been bred specifically for deep-sown seedling emergence. The BL training population was developed within the same breeding program compared to the DP that is composed of varieties from various breeding programs in the Pacific Northwest. As expected we see from both individual trials and across all scenarios, the BL had higher statistical mean accuracy than the DP. The training populations had similar accuracies for cross-validations, but the BLs displayed statistically higher accuracy for the independent validations as discussed previously. The increase in GS accuracy using the BL is advantageous for breeding programs because they can use their already existing breeding trials for use in GS without an expensive need to grow, screen, and genotype a diversity panel outside of their breeding program.

### GS Applications in Breeding for Complex Traits

Breeding for complex traits has until recently been implemented by creating genetic variation, selecting elite lines to cross, and turn the best progeny into improved cultivars which is dependent on phenotypic selection (Bernardo 2008). This traditional method is effective for seedling emergence. Even though the genetic architecture of seedling emergence remains elusive and depends on adequate field screening conditions and multiple factors, phenotypic selection has improved seedling emergence. In the dryland winter wheat region in Washington, where deep-sowing of winter wheat occurs, plant breeding efforts of 60 years have been able to develop modern genotypes that are fast emerging while having the GA insensitive mutant alleles *Rht-B1b* and *Rht-D1b*, and therefore, overcome the problems of historic genotypes. However, seedling emergence still relies on adequate environmental conditions for phenotypic screening and scoring. The dependence on the environment prevents phenotypic selection frequently as seen by the lack of BL trials from 2016 to 2019. The reliance on the environment creates an opportunity for constant genetic gain and selection using GS.

The dependence on the environment can be mitigated by appropriate replication and field design. However, the selection from the large unreplicated trials to the smaller replicated trials are an ideal stage to implement GS. GS will be more efficient in early generations trials within breeding programs that often use unreplicated trials in order to evaluate thousands of lines with limited seed material. A fraction of the large number of lines is then selected and screened in replicated yield trials. Therefore, we must account for the unreplicated nature of these trial designs and adequately account for it to help overcome environmental dependency. The lack of replications makes it difficult to separate genetic and non-genetic effects (Tolhurst et al. 2019). In the Washington State University breeding program, we use an ACBD design that allows the replication of check varieties and, therefore, enough degrees of freedom for estimating the experimental error and varietal effects (Federer and Raghavarao 1975). The justification of using adjusted means in our GS models can be seen by the lack of significant environmental effects in the ANOVA results caused by using a full ACBD model with genotype effects in combined trials. The lack of significance may be caused by convergence issues in the full models. Our results also showed the ability to overcome the environmental effect by combining trials within our training populations, and showing our ability to still accurately predict and potentially implement GS within early generation unreplicated trials.

The difficulties with the complex relationship between seedling emergence and the environment combined with unreplicated trials can be seen in the complex GS results. However, GS allows us to forgo traditional linkage mapping and GWAS studies based on an arbitrary threshold to conclude significant SNPs. Instead, GS allows us to use genome-wide markers that may account for genes that affect not just coleoptile length, but coleoptile diameter, emergence speed, emergence force, and lifting capacity of the first leaf. Even in the face of so many factors affecting seedling emergence, we were still able to achieve moderate accuracy in cross-validation schemes. Therefore, GS will be able to play an important role in selecting for deep-sown seedling emergence in winter wheat. In scenarios such as the BL populations from 2016-2019 where there were inadequate phenotypic variation and screening conditions, GS will be able to select for fast-emerging genotypes. Overall, the moderate accuracy of the genomic selection models will help breeders identify breeding lines with better seedling emergence in low rainfall environments and select seedling emergence even in years with little difference between breeding lines due to field conditions. The breeding lines selected will display superior seedling emergence and stand establishment, which will result in higher yield potential.

## Conclusion

This study displayed the ability for GS to predict a complex trait such as deep-sown seedling emergence. The comparison of GS models and the lack of significant differences between them indicated a trait with moderate heritability controlled by many loci. Both the complex nature of deep-sown seedling emergence and it’s relationship with the environment was indicated by the low validation set accuracy and the importance for further genetic dissection. This study opens the door for further investigation into the pleiotropic and environmental effects of a complex phenotypic trait. Even in the face of a complex trait, GS has still proven to have moderate accuracy and will enable breeders to select for a trait dependent on environmental conditions conducive for phenotypic selection. The environmental requirement is why genomic selection can play a helpful role in selection. Therefore, understanding the environmental effects on genomic prediction accuracy is crucial for adequately creating training populations and implementing GS for complex traits such as deep-sown seedling emergence in winter wheat. Overall, our study showed that breeders will be able to implement GS for a complex trait such as seedling emergence by using parametric models that model additive effects within their own breeding programs with increased accuracy as they combine phenotypic data over years. GS will allow breeders to increase genetic gain and build on the already 60 plus years of successful breeding for fast-emerging cultivars in a deep-sown low precipitation environment.

## Author’s contributions

LM: conceptualized the idea, analyzed data, and drafted the manuscript; AC: supervised the study, conducted field trials, edited the manuscript and obtained the funding for the project.

## Acknowledgments

The authors would like to acknowledge the Washington State University Winter Wheat Breeding Program personnel Gary Shelton for plot maintenance and data collection under field conditions. The authors would also like to acknowledge Brian P. Ward for sequence alignment, SNP calling, and quality filtering.

## Supplementary Material

Supplementary tables and images are available in a separate file.

## References

Allan RE, Vogel OA, Peterson CJ (1962) Seedling Emergence Rate of Fall-sown Wheat and Its Association with Plant Height and Coleoptile Length ^1^. Agron J 54:347–350. https://doi.org/10.2134/agronj1962.00021962005400040022x

Appels R, Eversole K, Feuillet C, et al (2018) Shifting the limits in wheat research and breeding using a fully annotated reference genome. Science 361:eaar7191

Arndt W (1965) The impedance of soil seals and the forces of emerging seedlings. Soil Res 3:55–68

Asoro FG, Newell MA, Beavis WD, et al (2011) Accuracy and Training Population Design for Genomic Selection on Quantitative Traits in Elite North American Oats. Plant Genome J 4:132. https://doi.org/10.3835/plantgenome2011.02.0007

Azodi CB, Bolger E, McCarren A, et al (2019) Benchmarking parametric and machine learning models for genomic prediction of complex traits. G3 Genes Genomes Genet 9:3691–3702

Balestre M, Von Pinho RG, Souza JC (2010) Prediction of maize single-cross performance by mixed linear models with microsatellite marker information. Genet Mol Res 9:1054–1068

Bernardo R (2008) Molecular markers and selection for complex traits in plants: Learning from the last 20 Years. Crop Sci 48:1649. https://doi.org/10.2135/cropsci2008.03.0131

Bernardo R (2014) Essentials of plant breeding. Stemma Press, Woodbury, Minnesota

Bernardo R (2010) Genomewide selection with minimal crossing in self-pollinated crops. Crop Sci 50:624. https://doi.org/10.2135/cropsci2009.05.0250

Bernardo R (2020) Breeding for quantitative traits in plants, 3rd ed. Stemma Press, Woodbury, Minnesota

Botwright T, Rebetzke G, Condon T, Richards R (2001) The effect of rht genotype and temperature on coleoptile growth and dry matter partitioning in young wheat seedlings. Funct Plant Biol 28:417–423. https://doi.org/10.1071/pp01010

Browning BL, Zhou Y, Browning SR (2018) A one-penny imputed genome from next-generation reference panels. Am J Hum Genet 103:338–348

Chastain TG, Ward KJ, Wysocki DJ (1995) Stand establishment responses of soft white winter wheat to seedbed residue and seed size. Crop Sci, 35:1

Crossa J, Pérez P, Hickey J, et al (2014) Genomic prediction in CIMMYT maize and wheat breeding programs. Heredity 112:48–60. https://doi.org/10.1038/hdy.2013.16

Cullis BR, Smith AB, Coombes NE (2006) On the design of early generation variety trials with correlated data. J Agric Biol Environ Stat 11:381. https://doi.org/10.1198/108571106X154443

Daetwyler HD, Pong-Wong R, Villanueva B, Woolliams JA (2010) The impact of genetic architecture on genome-wide evaluation methods. Genetics 185:1021–1031

de Mendiburu F, de Mendiburu MF (2019) Package ‘agricolae.’ R Package Version 1.2-8

Denis M, Bouvet J-M (2013) Efficiency of genomic selection with models including dominance effect in the context of Eucalyptus breeding. Tree Genet Genomes 9:37–51

Desta ZA, Ortiz R (2014) Genomic selection: genome-wide prediction in plant improvement. Trends Plant Sci 19:592–601. https://doi.org/10.1016/j.tplants.2014.05.006

Đorđević V, Ćeran M, Miladinović J, et al (2019) Exploring the performance of genomic prediction models for soybean yield using different validation approaches. Mol Breed 39:74

Dudley JW, Johnson GR (2009) Epistatic models improve prediction of performance in corn. Crop Sci 49:763–770

Elshire RJ, Glaubitz JC, Sun Q, et al (2011) A robust, simple genotyping-by-sequencing (GBS) approach for high diversity species. PLoS ONE 6:e19379. https://doi.org/10.1371/journal.pone.0019379

Endelman JB (2011) Ridge regression and other kernels for genomic selection with R package rrBLUP. Plant Genome J 4:250. https://doi.org/10.3835/plantgenome2011.08.0024

Federer WF (1956) Experimental design. LWW 81:4

Federer WT, Raghavarao D (1975) On augmented designs. Biometrics 31:29. https://doi.org/10.2307/2529707

Goldman I (2019) Plant breeding reviews. John Wiley & Sons

González-Camacho JM, Ornella L, Pérez-Rodríguez P, et al (2018) Applications of machine learning methods to genomic selection in breeding wheat for rust resistance. Plant Genome 11:0. https://doi.org/10.3835/plantgenome2017.11.0104

Habier D, Fernando RL, Dekkers JCM (2007) The impact of genetic relationship information on genome-assisted breeding values. Genetics 177:2389–2397. https://doi.org/10.1534/genetics.107.081190

Haile TA, Walkowiak S, N’Diaye A, et al (2020) Genomic prediction of agronomic traits in wheat using different models and cross-validation designs. Theor Appl Genet 1–18

Heffner EL, Jannink J-L, Iwata H, et al (2011a) Genomic selection accuracy for grain quality traits in biparental wheat populations. Crop Sci 51:2597. https://doi.org/10.2135/cropsci2011.05.0253

Heffner EL, Jannink J-L, Sorrells ME (2011b) Genomic selection accuracy using multifamily prediction models in a wheat breeding program. Plant Genome 4:65. https://doi.org/10.3835/plantgenome2010.12.0029

Heslot N, Rutkoski J, Poland J, et al (2013) Impact of marker ascertainment bias on genomic selection accuracy and estimates of genetic diversity. PLoS ONE 8:e74612. https://doi.org/10.1371/journal.pone.0074612

Heslot N, Yang H-P, Sorrells ME, Jannink J-L (2012) Genomic selection in plant breeding: A comparison of models. Crop Sci 52:146. https://doi.org/10.2135/cropsci2011.06.0297

Holland JB (2007) Genetic architecture of complex traits in plants. Curr Opin Plant Biol 10:156–161

Huang M, Ward B, Griffey C, et al (2018) The accuracy of genomic prediction between environments and populations for soft wheat traits. Crop Sci 58:2274. https://doi.org/10.2135/cropsci2017.10.0638

Jannink J-L, Lorenz AJ, Iwata H (2010) Genomic selection in plant breeding: from theory to practice. Brief Funct Genomics 9:166–177. https://doi.org/10.1093/bfgp/elq001

Jarquín D, Crossa J, Lacaze X, et al (2014) A reaction norm model for genomic selection using high-dimensional genomic and environmental data. Theor Appl Genet 127:595–607

Jia Z (2017) Controlling the overfitting of heritability in genomic selection through cross validation. Sci Rep 7:1–9

Juliana P, Singh RP, Singh PK, et al (2017) Comparison of models and whole-genome profiling approaches for genomic-enabled prediction of Septoria tritici blotch, Stagonospora nodorum blotch, and tan spot resistance in wheat. Plant Genome 10:1–16

Kamiak (2021) High Performance Computing | Washington State University. In: High Perform. Comput. https://hpc.wsu.edu/. Accessed 21 Jan 2021

Kuhn M (2008) Building predictive models in R using the caret package. J Stat Softw 28:1–26

Lande R, Thompson R (1990) Efficiency of marker-assisted selection in the improvement of quantitative traits. Genetics 124:743–756

Larkin, Lozada, Mason (2019) Genomic Selection—Considerations for successful implementation in wheat breeding programs. Agronomy 9:479. https://doi.org/10.3390/agronomy9090479

Li H, Durbin R (2009) Fast and accurate short read alignment with Burrows–Wheeler transform. bioinformatics 25:1754–1760

Lopes MS, Reynolds MP, Manes Y, et al (2012) Genetic yield gains and changes in associated traits of CIMMYT spring bread wheat in a “historic” set representing 30 years of breeding. Crop Sci 52:1123–1131

Lorenz AJ, Chao S, Asoro FG, et al (2011) Genomic selection in plant breeding: knowledge and prospects. Advances in Agronomy. Elsevier, pp 77–123

Lorenz AJ, Smith KP, Jannink J-L (2012) Potential and optimization of genomic selection for fusarium head blight resistance in six-row barley. Crop Sci 52:1609. https://doi.org/10.2135/cropsci2011.09.0503

Lozada DN, Carter AH (2019) Accuracy of single and multi-trait genomic prediction models for grain yield in US pacific northwest winter wheat. Crop Breed Genet Genomics. https://doi.org/10.20900/cbgg20190012

Lozada DN, Carter AH (2020) Insights into the genetic architecture of phenotypic stability traits in winter wheat. Agronomy 10:368

Lozada DN, Godoy JV, Ward BP, Carter AH (2019) Genomic prediction and indirect selection for grain yield in US pacific northwest winter wheat using spectral reflectance indices from high-throughput phenotyping. Int J Mol Sci 21:165. https://doi.org/10.3390/ijms21010165

Lozada DN, Ward BP, Carter AH (2020) Gains through selection for grain yield in a winter wheat breeding program. PLoS One 15:e0221603

Luo Z, Wang M, Long Y, et al (2017) Incorporating pleiotropic quantitative trait loci in dissection of complex traits: seed yield in rapeseed as an example. Theor Appl Genet 130:1569–1585

Lutcher LK, Wuest SB, Johlke TR (2019) First leaf emergence force of three deep-planted winter wheat cultivars. Crop Sci 59:772–777. https://doi.org/10.2135/cropsci2018.08.0495

Meuwissen THE, Hayes BJ, Goddard ME (2001) Prediction of total genetic value using genome-wide dense marker maps. Genetics 157:1819–1829

Michel S, Ametz C, Gungor H, et al (2016) Genomic selection across multiple breeding cycles in applied bread wheat breeding. Theor Appl Genet 129:1179–1189. https://doi.org/10.1007/s00122-016-2694-2

Mirdita V, He S, Zhao Y, et al (2015) Potential and limits of whole genome prediction of resistance to fusarium head blight and septoria tritici blotch in a vast central european elite winter wheat population. Theor Appl Genet 128:2471–2481

Mohan A, Schillinger WF, Gill KS (2013) Wheat seedling emergence from deep planting depths and its relationship with coleoptile length. PLoS ONE 8:e73314. https://doi.org/10.1371/journal.pone.0073314

Pérez P, de los Campos G (2014) Genome-wide regression and prediction with the BGLR statistical package. Genetics 198:483–495. https://doi.org/10.1534/genetics.114.164442

Pérez-Enciso M, Zingaretti LM (2019) A guide on deep learning for complex trait genomic prediction. Genes 10:553

Pérez-Rodríguez P, Gianola D, González-Camacho JM, et al (2012) Comparison Between Linear and Non-parametric Regression Models for Genome-Enabled Prediction in Wheat. G3amp58 GenesGenomesGenetics 2:1595–1605. https://doi.org/10.1534/g3.112.003665

Poland J, Endelman J, Dawson J, et al (2012) Genomic selection in wheat breeding using genotyping-by-sequencing. Plant Genome 5:103–113

R Core Team (2018) R: A language and environment for statistical computing. Version 3.5.1. R Foundation for Statistical Computing, Vienna, Austria. URL https://www.R-project.org/

Riedelsheimer C, Technow F, Melchinger AE (2012) Comparison of whole-genome prediction models for traits with contrasting genetic architecture in a diversity panel of maize inbred lines. BMC Genomics 13:452

Rolling WR, Dorrance AE, McHale LK (2020) Testing methods and statistical models of genomic prediction for quantitative disease resistance to Phytophthora sojae in soybean [Glycine max (L.) Merr] germplasm collections. Theor Appl Genet 133:3441–3454

Sandhu KS, Lozada DN, Zhang Z, et al (2020) Deep learning for predicting complex traits in spring wheat breeding program. Front Plant Sci 11:2084

SAS Institute, Inc (2011) SAS® 9.3 system options: Reference. SAS Institute Inc Cary, NC

Schillinger WF (2011) Rainfall impacts winter wheat seedling emergence from deep planting depths. Agron J 103:730–734. https://doi.org/10.2134/agronj2010.0442

Schillinger WF, Donaldson E, Allan RE, Jones SS (1998) Winter wheat seedling emergence from deep sowing depths. Agron J 90:582–586. https://doi.org/10.2134/agronj1998.00021962009000050002x

Schillinger WF, Schofstoll SE, Smith TA, Jacobsen JA (2017) Laboratory method to evaluate wheat seedling emergence from deep planting depths. Agron J 109:2004–2010. https://doi.org/10.2134/agronj2016.12.0715

Shikha M, Kanika A, Rao AR, et al (2017) Genomic selection for drought tolerance using genome-wide SNPs in maize. Front Plant Sci 8:550

Song F, Guo Z, Mei D (2010) Feature selection using principal component analysis. In: 2010 international conference on system science, engineering design and manufacturing informatization. IEEE, pp 27–30

Sunderman DW (1964) Seedling emergence of winter wheats and its association with depth of sowing, coleoptile length under various conditions, and plant height ^1^. Agron J 56:23–25. https://doi.org/10.2134/agronj1964.00021962005600010008x

Tang Y, Liu X, Wang J, et al (2016) GAPIT Version 2: An enhanced integrated tool for genomic association and prediction. Plant Genome 9:0. https://doi.org/10.3835/plantgenome2015.11.0120

Tolhurst DJ, Mathews KL, Smith AB, Cullis BR (2019) Genomic selection in multi-environment plant breeding trials using a factor analytic linear mixed model. J Anim Breed Genet 136:279–300. https://doi.org/10.1111/jbg.12404

Varona L, Legarra A, Toro MA, Vitezica ZG (2018) Non-additive effects in genomic selection. Front Genet 9:78. https://doi.org/10.3389/fgene.2018.00078

Wang J, Zhou Z, Zhang Z, et al (2018) Expanding the BLUP alphabet for genomic prediction adaptable to the genetic architectures of complex traits. Heredity 121:648–662. https://doi.org/10.1038/s41437-018-0075-0

Wang X, Yang Z, Xu C (2015) A comparison of genomic selection methods for breeding value prediction. Sci Bull 60:925–935

Ward BP, Brown-Guedira G, Tyagi P, et al (2019) Multienvironment and multitrait genomic selection models in unbalanced early-generation wheat yield trials. Crop Sci 59:491. https://doi.org/10.2135/cropsci2018.03.0189

Wellmann R, Bennewitz J (2012) Bayesian models with dominance effects for genomic evaluation of quantitative traits. Genet Res 94:21–37

Wickham H (2011) ggplot2. Wiley Interdiscip Rev Comput Stat 3:180–185

