## Supplementary Material for "Comparison of Genomic Selection Models for Exploring Predictive Ability of Complex Traits in Breeding Programs"

**Table S1.** Analysis of variance (ANOVA) for deep-sowing seedling emergence in individual and combined trials for the diversity panel (DP) training population and breeding line (BL) training population phenotyped from 2015 to 2019 and 2015 to 2020, respectively.

| Population | Set | Year | Check (Fixed/ Estimate) | Genotype (Random/ Variance) | Block (Random/ Variance) | Environment (Random/ Variance) | GEI (Random/ Variance) | Residual |
|---|---|---|---|---|---|---|---|---|
| DP | DP | 2015 | 17.21 | 161.90 * | 171.50 *** | | | 161.30 |
| DP | DP | 2017 | 3.56 | 45.28 * | 7.07 *** | | | 67.22 |
| DP | DP | 2018 | -0.05 | 149.76 | 0.00 | | | 50.00 |
| DP | DP | 2019 | -2.76 | 190.70 | 343.10 *** | | | 325.80 |
| DP | DP | 2015-2017 | 17.22 | 31.76 ** | 22.95 | 14.37 | 69.93 | 116.04 |
| DP | DP | 2015-2018 | 17.27 | 17.74 ** | 26.40 | 197.91 | 79.41 * | 114.77 |
| DP | DP | 2015-2019 | 17.34 | 25.36 *** | 25.05 | 43.27 | 118.83 * | 144.24 |
| BL | $F_{3:5}$ | 2015 | 11.22 | 164.80 *** | 109.40 *** | | | 142.50 |
| BL | DH | 2020 | -0.08 | 73.82 | 48.02 *** | | | 292.87 |
| BL | $F_{3:5}$ and DH | 2015-2020 | -0.01 | 115.70 | 128.50 | 128.80 | 117.80 | 123.70 |

0 '***' 0.001 '**' 0.01 '*'; DH: doubled-haploid; GEI: genotype-by-environment interaction

**Table S2.** Phenotypic correlations for deep-sowing seedling emergence adjusted means between individual and combined trials for the diversity panel (DP) training population and breeding line (BL) training population phenotyped from 2015 to 2019 and 2015 to 2020, respectively.

| | DP 2017 | DP 2018 | DP 2019 | DP 2015-2017 | DP 2015-2018 | DP 2015-2019 | $F_5$ 2015 | DH 2020 | $F_5$ 2015 and DH 2020 |
|---|---|---|---|---|---|---|---|---|---|
| DP 2015 | 0.50*** | 1.00 | 0.19* | 0.87*** | 0.94*** | 0.65*** | 0.93 | -0.95 | -0.27 |
| DP 2017 | | 0.50 | 0.94 | 0.87*** | 0.76*** | 0.98*** | 0.78* | -0.20 | 0.70 |
| DP 2018 | | | 0.19* | 0.87 | 0.94*** | 0.65*** | 0.93 | -0.95* | -0.27 |
| DP 2019 | | | | 0.65* | 0.50*** | 0.87*** | 0.54 | 0.14 | 0.90 |
| DP 2015-2017 | | | | | 0.98*** | 0.94*** | 0.99 | -0.66 | 0.25 |
| DP 2015-2018 | | | | | | 0.87*** | 1.00 | -0.79 | 0.06 |
| DP 2015-2019 | | | | | | | 0.89 | -0.38 | 0.55 |
| $F_{3:5}$ 2015 | | | | | | | | -0.76 | 0.11*** |
| DH 2020 | | | | | | | | | 0.56*** |
| $F_{3:5}$ 2015 and DH 2020 | | | | | | | | | |

0 '***' 0.001 '**' 0.01 '*'; DH: doubled-haploid

**Table S3.** Comparison of genomic selection model accuracy and pairwise comparisons for deep-sowing seedling emergence for Pacific Northwest winter wheat diversity panel (DP) lines phenotyped from 2015-2019 in Lind, WA using cross-validation within and across trials.

| Model | BayesA | BayesB | BayesC | BayesL | BayesRR | cBLUP | gBLUP | MK | RF_LD | RF_M | RF_PC | RKHS | rrBLUP | sBLUP | SVM_LD | SVM_M | SVM_PC |
|---|---|---|---|---|---|---|---|---|---|---|---|---|---|---|---|---|---|
| DP 2015 | 0.46(ab) | 0.46(a) | 0.45(abc) | 0.45(abc) | 0.45(bcd) | 0.45(abc) | 0.45(abc) | 0.44(de) | 0.45(bc) | 0.46(ab) | 0.3(g) | 0.44(cd) | 0.45(abc) | 0.46(a) | 0.43(e) | 0.44(de) | 0.34(f) |
| DP 2017 | 0.3(a) | 0.3(ab) | 0.29(abc) | 0.29(abcd) | 0.29(abc) | 0.28(bcd) | 0.3(a) | 0.28(d) | 0.24(e) | 0.25(e) | 0.14(g) | 0.28(cd) | 0.29(abcd) | 0.08(h) | 0.25(e) | 0.24(e) | 0.18(f) |
| DP 2018 | 0.45(a) | 0.45(a) | 0.45(a) | 0.45(a) | 0.45(a) | 0.45(a) | 0.45(a) | 0.45(a) | 0.45(a) | 0.44(a) | 0.35(c) | 0.45(a) | 0.45(a) | 0.18(d) | 0.45(a) | 0.42(b) | 0.36(c) |
| DP 2019 | 0.29(cdef) | 0.29(cdef) | 0.28(cdef) | 0.3(abc) | 0.28(ef) | 0.31(a) | 0.28(f) | 0.29(cde) | 0.31(ab) | 0.31(ab) | 0.22(g) | 0.3(bcd) | 0.28(def) | 0.16(h) | 0.29(bcd) | 0.29(cdef) | 0.23(g) |
| DP 2015-2017 | 0.46(a) | 0.45(a) | 0.45(ab) | 0.45(ab) | 0.45(ab) | 0.46(a) | 0.46(a) | 0.45(abc) | 0.44(abc | 0.43(cd) | 0.32(f) | 0.45(ab) | 0.45(a) | 0.44(abc | 0.43(bc) | 0.41(d) | 0.35€ |
| DP 2015-2018 | 0.5(cd) | 0.5(cd) | 0.49(cde) | 0.49(cde) | 0.49(cde) | 0.5(cd) | 0.5(cd) | 0.5(cd) | 0.51(b) | 0.52(a) | 0.43(g) | 0.5(bc) | 0.49(def) | 0.48(f) | 0.49(ef) | 0.49(ef) | 0.43(g) |
| DP 2015-2019 | 0.48(cde) | 0.48(cde) | 0.48(cde) | 0.48(cde) | 0.48(def) | 0.49(ab) | 0.48(cde) | 0.48(cd) | 0.5(ab) | 0.5(a) | 0.42(h) | 0.49(bc) | 0.48(def) | 0.44(g) | 0.47(ef) | 0.47(f) | 0.42(h) |
| Overall | 0.42(a) | 0.42(a) | 0.41(ab) | 0.42(ab) | 0.41(ab) | 0.42(a) | 0.42(ab) | 0.41(ab) | 0.41(ab) | 0.41(ab) | 0.31(c) | 0.42(ab) | 0.41(ab) | 0.32(c) | 0.4(ab) | 0.39(b) | 0.33(c) |

Models labeled with the same letter are not significantly different (P-value =0.05)

**Table S4.** Comparison of genomic selection model accuracy and pairwise comparisons for deep-sowing seedling emergence in a Pacific Northwest winter wheat breeding lines (BL) phenotyped in 2015-2020 in Lind, WA using cross-validation within and across trials.

| Model | BayesA | BayesB | BayesC | BayesL | BayesRR | cBLUP | gBLUP | MK | RF_LD | RF_M | RF_PC | RKHS | rrBLUP | sBLUP | SVM_LD | SVM_M | SVM_PC |
|---|---|---|---|---|---|---|---|---|---|---|---|---|---|---|---|---|---|
| $F_{3:5}$ 2015 | 0.54(ab) | 0.54(ab) | 0.54(ab) | 0.54(ab) | 0.54(ab) | 0.55(ab) | 0.54(b) | 0.55(ab) | 0.55(ab) | 0.55(ab) | 0.38(c) | 0.55(ab) | 0.54(ab) | 0.34(d) | 0.55(ab) | 0.56(a) | -0.06(e) |
| DH 2020 | 0.41(ab) | 0.4(abc) | 0.4(abc) | 0.4(abc) | 0.4(abc) | 0.4(abc) | 0.4(abc) | 0.4(abc) | 0.39(bc) | 0.4(abc) | 0.32(d) | 0.41(a) | 0.41(a) | 0.25(e) | 0.39(c) | 0.41(a) | -0.07(f) |
| $F_{3:5}$ 2015 and DH 2020 | 0.43(bcd) | 0.43(bcd) | 0.43(bcd) | 0.43(bcd) | 0.43(bcd) | 0.44(ab) | 0.43(abcd) | 0.44(ab) | 0.42(d) | 0.42(cd) | 0.35(e) | 0.44(a) | 0.43(abc) | 0.22(f) | 0.43(abc) | 0.42(cd) | -0.03(g) |
| Overall | 0.46(a) | 0.46(a) | 0.46(a) | 0.46(a) | 0.46(a) | 0.46(a) | 0.46(a) | 0.46(a) | 0.45(a) | 0.46(a) | 0.35(b) | 0.47(a) | 0.46(a) | 0.27(c) | 0.46(a) | 0.46(a) | -0.06(d) |

Models labeled with the same letter are not significantly different (P-value =0.05); DH: Doubled-Haploid

**Table S5.** Comparison of genomic selection model accuracy and pairwise comparisons for deep-sowing seedling emergence in a Pacific Northwest winter wheat diversity panel (DP) phenotyped from 2015-2019 in Lind, WA for independent validations using independent validations of both the DP and breeding line (BL) training populations.

| Model | BayesA | BayesB | BayesC | BayesL | BayesRR | cBLUP | gBLUP | MK | RF_LD | RF_M | RF_PC | RKHS | rrBLUP | sBLUP | SVM_LD | SVM_M | SVM_PC |
|---|---|---|---|---|---|---|---|---|---|---|---|---|---|---|---|---|---|
| DP 2015 to DP 2017 | 0.2(de) | 0.2(ef) | 0.21(bcd) | 0.2(de) | 0.21(b) | 0.21(bcd) | 0.21(bcde) | 0.21(bcde) | 0.2(g) | 0.2(fg) | 0.22(a) | 0.21(bc) | 0.21(cde) | 0.19(h) | 0.2(fg) | 0.19(h) | 0.13(i) |
| DP 2017 to DP 2018 | 0.07(cdef) | 0.07(bcd) | 0.07(abc) | 0.07(efgh) | 0.08(a) | 0.06(h) | 0.07(defg) | 0.07(bcd) | 0.06(h) | 0.06(h) | 0.07(cde) | 0.08(ab) | 0.07(fgh) | 0.07(cde) | 0.07(cd) | 0.07(gh) | 0.07(cde) |
| DP 2018 to DP 2019 | 0.11(ef) | 0.11(def) | 0.11(def) | 0.11(bcdef) | 0.11(f) | 0.12(abcde) | 0.11(bcdef) | 0.12(a) | 0.12(abc) | 0.12(ab) | 0.12(a) | 0.12(a) | 0.11(cdef) | 0.05(h) | 0.11(f) | 0.12(abcd) | 0.1(g) |
| DP 2015-2017 to DP 2018 | 0.06(ef) | 0.06(d) | 0.06(de) | 0.05(g) | 0.07(c) | 0.08(b) | 0.05(f) | 0.03(i) | 0.02(k) | 0.02(j) | 0.04(h) | 0.04(h) | 0.05(g) | 0.11(a) | 0.02(l) | 0.01(m) | 0(n) |
| DP 2015-2018 to DP 2019 | 0.18(abc) | 0.18(bcd) | 0.18(ab) | 0.17(cd) | 0.18(a) | 0.16(fg) | 0.17(d) | 0.17(fg) | 0.17(ef) | 0.17(efg) | 0.17(d) | 0.17(e) | 0.17(ef) | 0.17(ef) | 0.16(h) | 0.16(i) | 0.16(gh) |
| DP 2015-2019 to $F_5$ 2015 | 0.01(d) | 0.01(de) | 0.01(de) | 0(ef) | 0.02(d) | 0(f) | 0.02(d) | 0.04(c) | 0.01(de) | 0.03(c) | 0.08(b) | 0.04(c) | 0.04(c) | 0.11(a) | -0.01(g) | -0.02(h) | -0.08(i) |
| DP 2015-2019 to DH 2020 | 0.19(ab) | 0.2(ab) | 0.2(ab) | 0.19(ab) | 0.21(a) | -0.19(i) | 0.18(b) | 0.21(a) | -0.1(g) | -0.03(f) | 0.05(e) | 0.2(ab) | 0.16(c) | 0.1(d) | -0.14(h) | -0.18(i) | 0.19(ab) |
| Overall | 0.12(a) | 0.12(a) | 0.12(a) | 0.11(a) | 0.12(a) | 0.06(bc) | 0.12(a) | 0.12(a) | 0.07(bc) | 0.08(b) | 0.11(a) | 0.12(a) | 0.11(a) | 0.11(a) | 0.06(c) | 0.05(c) | 0.08(b) |

Models labeled with the same letter are not significantly different (P-value =0.05); DH: Doubled-Haploid

**Table S6.** Comparison of genomic selection model accuracy and pairwise comparisons for deep-sowing seedling emergence in a Pacific Northwest winter wheat breeding lines (BL) phenotyped in 2015-2020 in Lind, WA for independent validations of both the diversity panel (DP) and BL training populations.

| Model | BayesA | BayesB | BayesC | BayesL | BayesRR | cBLUP | gBLUP | MK | RF_LD | RF_M | RF_PC | RKHS | rrBLUP | sBLUP | SVM_LD | SVM_M | SVM_PC |
|---|---|---|---|---|---|---|---|---|---|---|---|---|---|---|---|---|---|
| $F_{3:5}$ 2015 to DH 2020 | -0.14(cd) | -0.15(cd) | -0.15(cd) | -0.14(cd) | -0.15(cd) | -0.28(g) | -0.17(de) | -0.12(c) | -0.18(de) | -0.12(c) | -0.07(b) | -0.16(cd) | -0.16(cd) | 0.13(a) | -0.24(fg) | -0.21(ef) | -0.12(c) |
| $F_{3:5}$ 2015 and DH 2020 to DP 2015 | 0.3(abc) | 0.3(abc) | 0.29(c) | 0.29(bc) | 0.3(abc) | 0.32(abc) | 0.31(abc) | 0.22(ef) | 0.32(a) | 0.32(ab) | 0.17(g) | 0.23(def) | 0.3(abc) | 0.26(d) | 0.22(f) | 0.24(de) | 0.1(h) |
| $F_{3:5}$ 2015 and DH 2020 to DP 2017 | 0.3(ab) | 0.3(ab) | 0.29(b) | 0.29(b) | 0.3(ab) | 0.32(ab) | 0.31(ab) | 0.22(de) | 0.13(g) | 0.32(a) | 0.17(f) | 0.23(cde) | 0.3(ab) | 0.26(c) | 0.22(e) | 0.24(cd) | 0.1(g) |
| $F_{3:5}$ 2015 and DH 2020 to DP 2018 | 0.3(ab) | 0.3(ab) | 0.29(b) | 0.29(b) | 0.3(ab) | 0.32(ab) | 0.31(ab) | 0.22(de) | -0.07(h) | 0.32(a) | 0.17(f) | 0.23(cde) | 0.3(ab) | 0.26(c) | 0.22(e) | 0.24(cd) | 0.1(g) |
| $F_{3:5}$ 2015 and DH 2020 to DP 2018 | 0.3(ab) | 0.3(ab) | 0.29(b) | 0.29(b) | 0.3(ab) | 0.32(ab) | 0.31(ab) | 0.22(de) | 0.08(g) | 0.32(a) | 0.17(f) | 0.23(cde) | 0.3(ab) | 0.26(c) | 0.22(e) | 0.24(cd) | 0.1(g) |
| Overall | 0.21(a) | 0.21(a) | 0.2(ab) | 0.2(ab) | 0.21(a) | 0.2(ab) | 0.21(a) | 0.15(bc) | 0.06(d) | 0.23(a) | 0.13(c) | 0.15(bc) | 0.21(a) | 0.23(a) | 0.13(c) | 0.15(bc) | 0.06(d) |

Models labeled with the same letter are not significantly different (P-value =0.05); DH: Doubled-Haploid

**Table S7.** Comparison of genomic selection scenarios for deep-sowing seedling emergence for cross-validations (CV) and both types of independent validations (IV), continuous training (CT) and validation sets (VS), in a Pacific Northwest winter wheat diversity panel (DP) and breeding lines (BL) phenotyped in 2015-2020 in Lind, WA.

| DP CV | | BL CV | | DP IV | | BL IV | |
|---|---|---|---|---|---|---|---|
| Scenario | Accuracy | Scenario | Accuracy | Scenario | Accuracy | Scenario | Accuracy |
| DP 2015 (CV) | 0.43(c) | $F_{3:5}$ 2015 (CV) | 0.49(a) | DP 2015 to DP 2017 (CT) | 0.2(a) | $F_{3:5}$ 2015 to DH 2020 (CT) | -0.14(c) |
| DP 2017 (CV) | 0.25(f) | DH 2020 (CV) | 0.36(c) | DP 2017 to DP 2018 (CT) | 0.07(e) | $F_{3:5}$ and DH to DP 2015 (VS) | 0.26(a) |
| DP 2018 (CV) | 0.42(d) | $F_{3:5}$ 2015 and DH 2020 (CV) | 0.39(b) | DP 2018 to DP 2019 (CT) | 0.11(c) | $F_{3:5}$ and DH to DP 2017 (VS) | 0.25(b) |
| DP 2019 (CV) | 0.28(e) | | | DP 2015-2017 to DP 2018 (CT) | 0.05(f) | $F_{3:5}$ and DH to DP 2018 (VS) | 0.24(b) |
| DP 2015-2017 (CV) | 0.43(c) | | | DP 2015-2018 to DP 2019 (CT) | 0.17(b) | $F_{3:5}$ and DH to DP 2018 (VS) | 0.25(b) |
| DP 2015-2018 (CV) | 0.49(a) | | | DP 2015-2019 to $F_{3:5}$ 2015 (VS) | 0.02(g) | | |
| DP 2015-2019 (CV) | 0.47(b) | | | DP 2015-2019 to DH 2020 (VS) | 0.09(d) | | |

Models labeled with the same letter are not significantly different (P-value =0.05); DH: Doubled-Haploid
